# Social Rank Alters the Synchrony of Attribute Integration in Altruistic Decisions

**DOI:** 10.1101/2023.10.26.564311

**Authors:** Yinmei Ni, Jian Li

## Abstract

Social rank, which represents the hierarchical dominance structure in societies, forms the backdrop against which most social decisions are made. Effective social decision-making demands the flexible integration of multifaceted information, including the relative weights and timings of different attributes. However, there is limited understanding of the potential impact of attribute timing on social preferences and, crucially, whether attribute weights and timings are susceptible to the influence of social rank. Here, through a two-stage social decision task, we manipulated subjects’ social rank before they engaged in altruistic decisions. By interrogating behavioral data through the lens of the time-varying drift diffusion process, we found that the behavioral patterns are better explained by a model that considers both attribute timing and weight. Interestingly, varying social ranks only modulate attribute timings in the decision process, leaving attribute weights unaffected. Furthermore, we demonstrated that individuals with more prosocial tendencies exhibited higher sensitivities of attribute timing in response to the changes of social rank and these results were replicated in a separate and larger cohort of participants. Our results underscore the intricate interplay between social rank and social attribute integration and introduce a new dimension to the computational mechanisms that underlie social decision-making.

**Author Summary:** Social rank characterizes the background where most social decisions take place, however, the cognitive and computational mechanisms underlying how social rank influences altruistic decisions remain elusive. Here, combining a two-stage social decision task and time-varying DDM, we reveal that social rank specifically modulates attribute timings in the decision process, leaving attribute weights unaffected. Our results underscore the intricate interplay between social rank and social attribute integration during social decision-making.

## Introduction

Social decisions are rarely made in a vacuum. The prevalence of social rank, the hierarchical dominance structure in the societies characterizes the background where most social decisions take place [1]. Adam Smith noted that people were willing to expend effort to attain certain social rank, which was thought to be associated with the expectation of entitlement to resources [2]. Indeed, social rank affects the allocation of resources. For instance, sellers or buyers with higher social rank in a market reap a greater share of the market surplus than their lower-rank counterparts [1]. Despite the pervasiveness of social ranking in any society, however, the cognitive and computational mechanisms underlying how social rank influences social decisions remain elusive.

According to formal decision theories, prosocial decision-making involves the integration of multiple attributes in the decision process [3–12]. A well-established fact is that individuals often weigh their own and others’ welfare when faced with resource allocation decisions. A variety of utility models have been proposed by emphasizing different components of the cost-benefit analysis, such as those focusing on the attributes of self-interest and inequity [6], selfishness and altruism [3, 11], or equity and efficiency [5,13]. These models assume that different attributes, combined with their subjective weights, enter into an evidence accumulation process, thus capturing the choice and response time patterns in altruistic decision-making [4,8]. However, such static models, in which the relative impact of each attribute remains constant during the decision process, are agnostic to the empirical findings that subjects’ social preference may change as a function of response time [14–16].

To address this question, recent work in social psychology has introduced dual-process models which propose that decisions evolve through continuous arbitration between fast, automatic dispositional preferences and slower, reflective and deliberate preferences during the decision process [14,16,17]. Nonetheless, both correlational and causal evidence suggests that the preference for more prosocial choices is not monotonically associated with response time [18–21], raising the question of whether decision conflict, rather than automatic or deliberate preferences, influences choice response time in social dilemmas [20,21]. Other lines of research, instead, adopted more algorithmically dynamic models, either by imposing an initial bias term to account for subjects’ dispositional preference in the evidence accumulation process [4], or by deploying differential subjective attribute weights according to participants’ gaze-informed attentional preference, to explain the dynamic evolvement of social preference [12]. Finally, evidence from individual food choice studies suggest that the temporal order in which different decision attributes are integrated in decision-making also affects choice preference and RT [22,23]. That is, both the symmetry (attribute weight) and synchrony (attribute timing) of attributes integration are likely to shape social preference across individuals and decision contexts. Delineating the channels via which social rank exerts its influence in resource allocation will pave the way to better understand how social preference can be flexibly modulated by specific social decision context.

To formally investigate the computational mechanism of how social rank affects social preference, here we propose an asynchronous multi-attribute integration model for altruistic choices. This model builds upon the canonical drift diffusion model (DDM), different variants of which are widely used in individual perceptual and value-based economic decision making [24–30]. Specifically, we adopted the multi-attribute, time-dependent version of the DDM (mtDDM) [22,23, 31,32]. The mtDDM was originally developed to capture the dynamic titration between tastiness and healthiness attributes during food selection decisions, but also has the potential to extend to altruistic choices where both self-interest and social factors (e.g., the welfare of others and the relative social rank) serve as driving attributes for subjects’ prosocial behavior.

Therefore, such a model allows us to ascribe subjects’ choice and response time (RT) data to two separate and well-defined psychological aspects in altruistic decisions: attribute timing, the time of certain decision attribute entering into the decision process, and attribute weight, the relative importance of specific attributes influencing choice selection. Previous studies have shown that both attribute timing and weight could bias subjects’ preferences and RTs. For example, it was demonstrated that dynamic visual attention shaped how strongly certain option or aspects (such as tastiness or healthiness) influence food preference using eye tracking and computer mouse tracking methods [25,32] and the revelation of food calorie information sped up the integration of healthiness attribute in the overweight individuals in a food-choice task [33]. Similarly, recently application of DDM models incorporating eye-gaze informed attribute weight to social decision-making studies have shed light on the computational and neural mechanisms underlying value integration and preference generation in altruistic decisions [8,12].

Moreover, by focusing on attribute timing and weight, the mtDDM avoids the debates surrounding the dual-process models, which hypothesize social decisions result from competing preferences of automatic disposition and controlled reflection. Instead, the model proposes that dynamic social preferences arise from the combined influence of attribute weight and timing during the decision process where attribute weights are associated with subjects’ dispositional social preference [34]. Importantly, we hypothesize that the temporal order of taking each attribute into consideration during the decision process (attribute timing) also plays a key role in shaping participants’ social preference and social rank may influence the synchrony of attribute integration. Indeed, previous studies have shown that in a money redistribution task with assigned social ranks, subjects’ social preference were heavily influenced by their initial social ranks, highlighting the intricate link between social rank and social preference [35,36]. This hypothesis is also consistent with research showing that the orders of queries about choice alternatives have a causal effect in decision preference [37]. For example, the endowment effect can emerge from different query orders between buyers and sellers [38]. Under this framework, therefore, flexible social decisions can be driven by the idiosyncratic attribute weights, attribute timing, or both. Moreover, attribute timing may be susceptible to the decision context and the endowment of different social rank may reveal the asynchrony of attribute integration.

We set out to test whether attribute timing and weight have distinctive contributions to social decision and how they are modulated by the corresponding social rank. Specifically, we designed a two-stage decision task where different social ranks were experimentally elicited before subjects completed an altruistic choice task (Fig.1). The model-based analyses (mtDDM) reveal that the timing and weights of two attributes, namely subjects’ own and others’ payoffs play dissociable and significant roles in the altruistic decision process. Interestingly, higher social rank leads to faster consideration of one’s own payoff relative to that of the other’s, but does not influence how strongly subjects evaluate their own or others’ payoffs. Furthermore, we show that the sensitivity of attribute timing in response to the change of social rank is mainly driven by the prosocial group in the population. The individualistic group, however, has a significantly higher weight towards their own payoff than the prosocial group but their attribute timing is less sensitive to the change of social rank than the prosocial subjects. Finally, these results were further replicated in a larger and separate cohort of participants, confirming our hypothesis that social rank affects social decisions via the modulation of attribute timing as opposed to the attribute weight.

**Fig 1.**
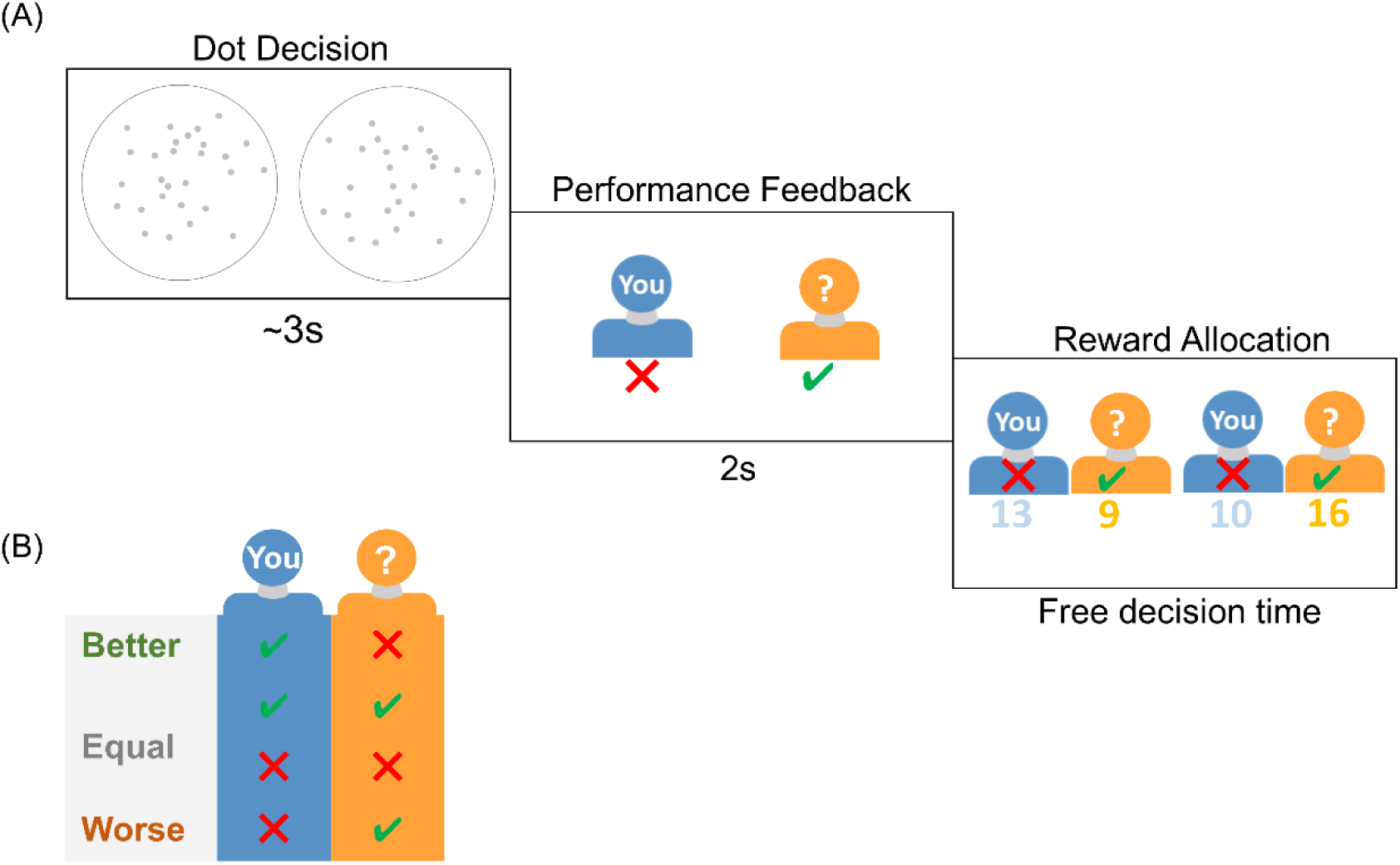
Task design. (A) Trial timeline. In each trial subjects first performed a dot number estimation task. Upon receiving feedbacks, they had to choose between two reward allocation options. (B) Four types of feedbacks were sorted into better, equal, and worse social rank conditions (see results) defined as the relative performance compared to their co-players.

## Results

### Social rank modulates individual social preference

In our study, trial by trial subjects were asked to make a reward allocation choice after the dot number estimation choice task paired with an anonymous co-player (Fig 1A). Based on the specific feedbacks from the dot estimation task (Fig 1B), we grouped trials into three social rank categories (better, equal, and worse, see methods and S1 Fig for details) as the relative performance of the subjects compared to their co-players and hence a proxy of subjects’ relative social rank in each trial (Fig 1B).

First, we examined how subjects’ social rank influenced their prosocial behavior and found that subjects’ self-payoff (*M_s_*) maximization behavior declined as their social rank worsened, whereas subjects were more likely to choose options that maximized their co-players’ payoffs (*M_o_*) (Fig 2A). One-way repeated ANOVA confirmed that the main effect of social rank was significant in both choice types (self-payoff maximization *M_s_*: F(2,128) = 23.167, P < 0.001; other-payoff maximization*M_o_*: F(2,128) = 20.649, P < 0.001). Next, we conducted a mixed-effect logistic regression analysis to quantitatively examine the effect of social rank on subjects’ social preferences by modeling subjects’ choices as a function of the difference of *M_s_* (Δ*M_s_*) and *M_o_* (Δ*M_o_*) between alternative options. Our results showed that the decision betas (regression coefficients) of Δ*M_s_* and Δ*M_o_* were both positive and significant across social rank (all Ps < 0.001), indicating that subjects evaluated both their own payoff as well as co-players’ during decisions. Consistent with previous findings (Fig 2A), social rank also exerted opposite effects on Δ*M_s_* and Δ*M_o_*: repeated one-way ANOVA showed that the influence of Δ*M_s_* declined with worsening social rank (Δ*M_s_*: F(2,128) = 215.045, P < 0.001, Fig 2B), whereas the effect of Δ*M_o_* increased (Δ*M_o_*: F(2,128) = 15.735, P < 0.001, Fig 2C), suggesting that subjects became more prosocial as their social rank decreased. Finally, the mean response time (RT) increased significantly as social rank worsened (Fig 2D, log RT: F(2,128) = 10.277, P < 0.001). These results on the surface support the hypothesis that social rank modulates decision weights (decision betas) on different decision attributes (Δ*M_s_* and Δ*M_o_*). Given the larger magnitude of the decision weight on Δ*M_s_* relative to that of Δ*M_o_*, worsening social rank, according to the standard drift diffusion model (DDM), would lead to more commensurate attribute weights of Δ*M_s_* and Δ*M_o_* and therefore prolong the decision process with longer RT. However, recent development in evidence accumulation models suggests that the rate of attribute accumulation and the time attributes enter into the decision process can have dissociable impacts on subjects’ choice and RT [22,23,31].

**Fig 2.**
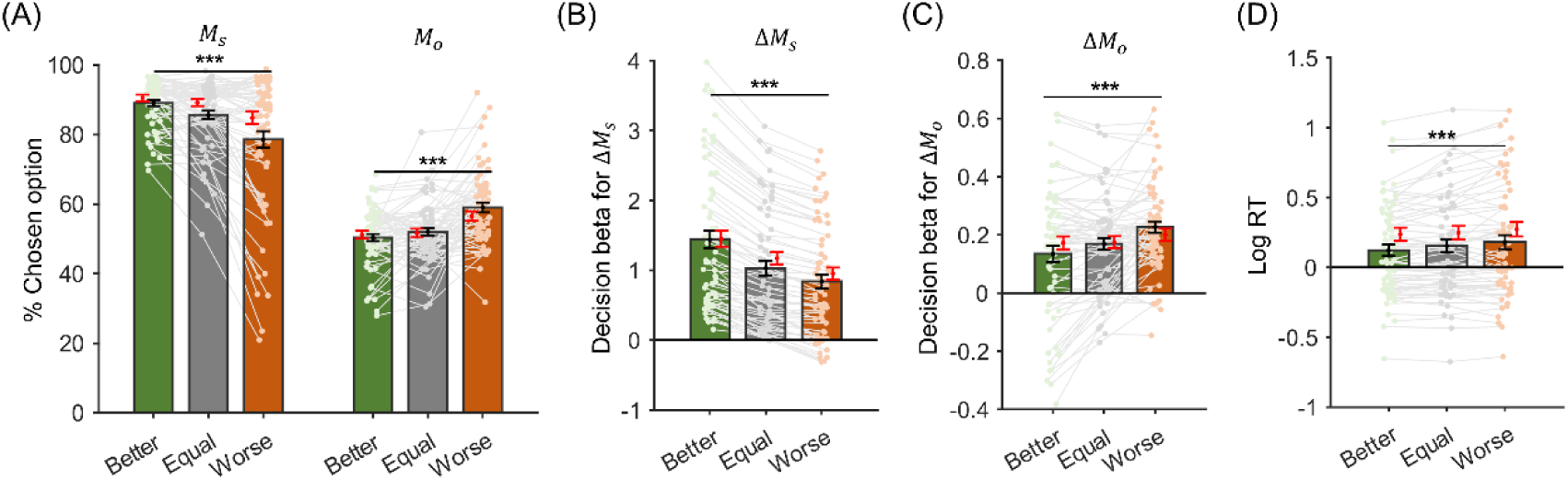
Behavioral results. (A) Percentage of choosing larger *M_s_* or larger *M_o_* options *M_s_*: The proportion of chosen option with larger *M_s_*; *M_o_*: the proportion of chosen option with larger *M_o_*. (B-C), Decision betas of Δ*M_s_* (B) and Δ*M_o_*(C) for choices across three social rank conditions. (d) Mean log response time (RT). Black error bars represent s.e.m across subjects and red error bars represent model predictions from the cross-validation results (see methods). *** p < 0.001.

### Time-varying drift diffusion process modeling

To systematically investigate how social rank affects subjects’ prosocial preferences, we adopted a sequential sampling model in which both the timing and weighting strength of different decision attributes (Δ*M_s_* and Δ*M_o_* in our task) are susceptible to the social rank. The timing or the temporal onsets of different attributes entering the decision process effectively controls the temporal dynamics of attribute weighting strength during the choice. Similar to earlier research developing such a framework [22,23,29,30,39–43], the multi-attribute time-varying drift diffusion model (mtDDM) is achieved by extending the classic DDM with an additional free parameter describing how fast one decision attribute, relative to the other, begins to be considered in the evidence accumulation process (see methods for details). Under this framework, the relative start time (RST) parameter, in combination with attribute weight, determines subjects’ prosocial preference and RT distribution. Figure 3A provides an example where Δ*M_s_* enters into the decision process earlier than Δ*M_o_*. According to the mtDDM, choices made before Δ*M_o_* is taken into account are predominantly influenced by the evidence accumulation rate of Δ*M_s_* (blue traces). On the contrary, Δ*M_o_* dominates choice selection before Δ*M_s_* enters into the decision process (Fig 3B, orange traces). Such effects are further reflected on the dynamic decision betas (logistic regression coefficients) of both Δ*M_s_* and Δ*M_o_* on subjects’ choice behavior as a function of RT (Fig 3C-E). More specifically, time-varying decision betas are observed when Δ*M_s_* starts in the decision process earlier (Fig 3C, orange trace Δ*M_o_*), or later (Fig 3E, blue trace Δ*M_s_*) than Δ*M_o_*. Such variability is caused by the asynchrony of Δ*M_s_* and Δ*M_o_* in terms of their temporal onsets in influencing the evidence accumulation process. Indeed, simultaneous consideration of both Δ*M_s_* and Δ*M_o_* yields relatively stable decision betas for different attributes (Δ*M_s_* and Δ*M_o_*) during the decision process (Fig 3D).

**Fig 3.**
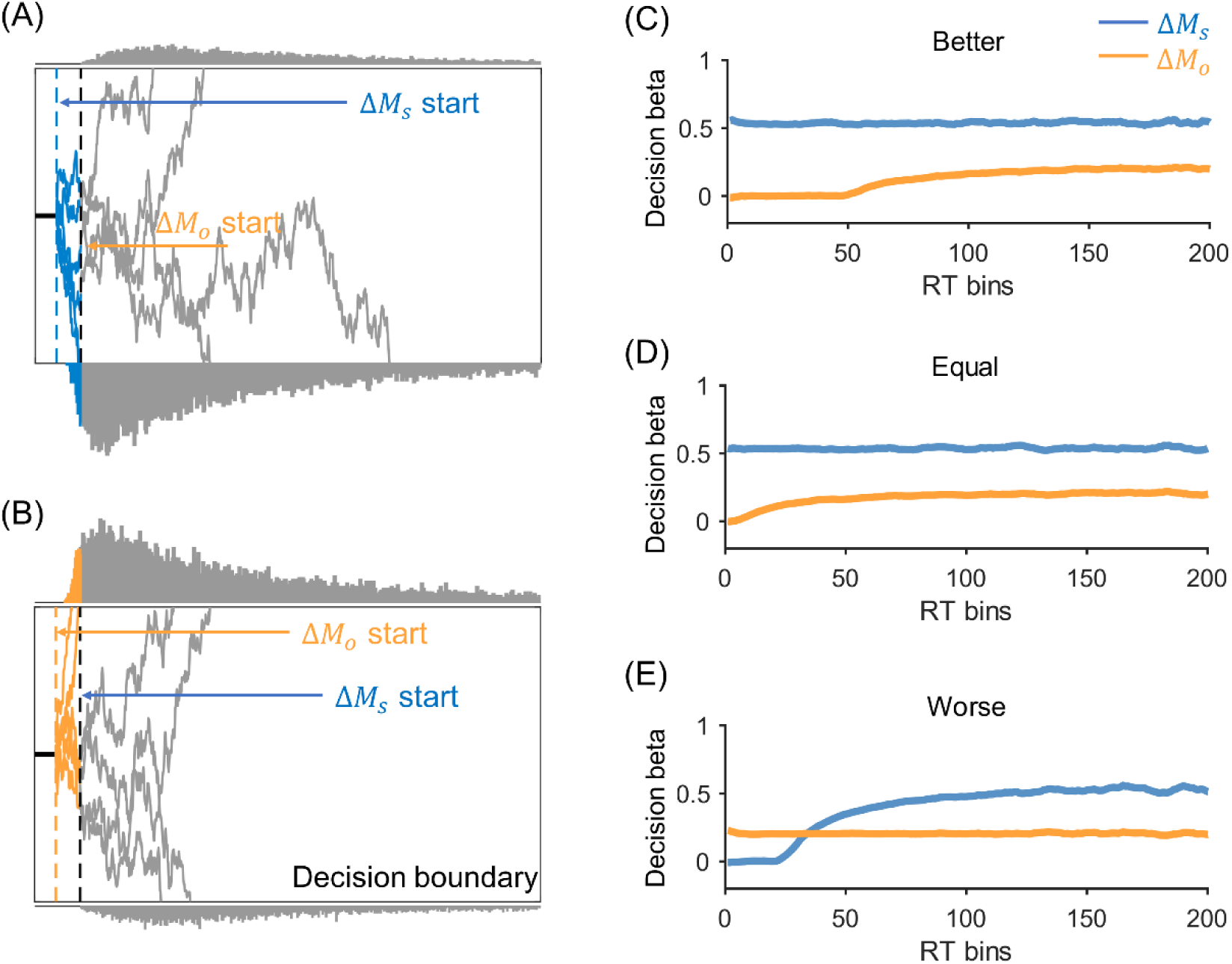
Model illustration and prediction. (A-B) are graphic illustrations of the time-varying drift diffusion process (mtDDM). (A) Illustration of when Δ*M_s_* enters the decision process first followed by Δ*M_o_*, and their start time difference is the relative start time (RST). A decision is made when the accumulated evidence reaches the decision boundary. Here we show that two agents with the same drift weight for the two attributes but different attribute timing onsets (Δ*M_s_* earlier in panel (A) and Δ*M_o_* earlier in panel (B)), could result in different choices. Blue, evidence accumulation trajectory where only Δ*M_s_* is considered in the decision process; orange, only Δ*M_o_* is considered; grey, both attributes accumulate in the decision process. (C), (D) and (E) are mtDDM simulated attribute effects (decision betas) as a function of response time (RT) for both Δ*M_s_* (blue) and Δ*M_o_* (yellow) in better, equal, and worse social conditions. The x-axis represents overlapping 100ms RT windows that slide along RT in steps of 10ms.

Recently, this approach has been applied to studies of individual preference formation involving the trade-off between attributes such as food tastiness and healthiness [23]. We adopted this framework to investigate whether, and if so, how the relative social standing would influence the timing of social information entering into the decision process and subsequently give rise to subjects’ prosocial preference. Based on the behavioral effects of social standings (Fig 2), three hypotheses related to the influence of social rank arise: (1) Social rank modulates both the accumulation rate (drift weight) and the time onsets of different attributes (Δ*M_s_* and Δ*M_o_*); (2) it only modulates the relative start time (RST) of attributes but not the drift weight and (3) it specifically modulates drift weight whereas has no effect on RST. To test these different possibilities, we built a series of multi-attribute time-varying DDMs (mtDDM) by including a relative start time (RST: the time onset difference between Δ*M_s_* and Δ*M_o_*) parameter in the standard drift diffusion model. In model 1, we assume that both Δ*M_s_* and Δ*M_o_* enter into the decision process simultaneously (RST = 0) and social rank modulates the individual drift weights of Δ*M_s_* and Δ*M_o_* respectively (Model1, see methods, Eq. 1). Furthermore, RST might be different across social rank and Δ*M_s_* and Δ*M_o_* share the same drift weight (Model 2). Finally, it is also possible that social rank modulates the individual drift weights Δ*M_s_* and Δ*M_o_* separately but with a constant RST (Model 3) or rank-related RSTs (Model 4). All the above models, in principle, can produce qualitatively similar behavioral patterns such as rank-dependent reaction time and decision betas shown above (Fig 2).

To quantitatively examine the resolution of the decision process, we used the differential evolution algorithm [44] (see methods for detailed description) to interrogate the candidate models against both behavioral choices reaction times of subjects’ data. This algorithm has been applied previously in both social and non-social preference tasks and is capable of distinguishing the timing and strength of different attributes influencing choices [22,45]. We compared our candidate models by performing a Bayesian model comparison [46] test and the model with social rank-dependent RSTs but rank-independent drift weights outperformed all other models (Model2: exceedance probability (EXP) = 0.999, S2A Fig). Further parameter recovery analysis showed that all the model parameters recovered well from simulated choice dataset with sufficient true parameter ranges of our winning model (RST model; S3 Fig).

We then conducted a series of analyses to test whether the RST model captured the characteristics of behavioral patterns of our subjects. First, we ran a cross-validation analysis to test the out-of-sample accuracy of our model in terms of predicting prosocial choices as well as RTs. We estimated the best-fitting parameters of the model from a randomly selected half of the trials (for each social rank) for each subject and tested the fit between model-predictions and observed data on the other half dataset (see methods for details). Model predictions captured the overall RT patterns in different social rank (Fig 2D, one-way repeated ANOVA, F(2,128) = 7.773, P < 0.001, red error bars) as well as the inter-subject difference within each social rank (S4 Fig). Furthermore, our model also captured subjects’ preference towards more prosocial options as their social rank decreased (Fig 2A, larger *M_s_* option proportion: F(2,128) = 14.183, P < 0.001; larger *M_o_* option proportion: F(2,128) = 12.258, P < 0.001, red error bars). The regression analysis also confirmed this pattern: Simulated choice data from our RST model revealed rank-dependent decision weights on both Δ*M_s_* (Fig 2B, F(2,128) = 205.167, P < 0.001, red error bars) and Δ*M_o_* (Fig 2C, F(2,128) = 6.555, P = 0.002, red error bars), suggesting the asynchrony of different decision attributes (RST) can sufficiently account for the choice and response time patterns we observed.

### Model parameters and inter-subject behavioral variance

Further examination of the best fitting model (the RST model) suggests that higher drift weights are assigned to the private attribute (Δ*M_s_*) than the social one (Δ*M_o_*) and this asymmetry does not change across social rank (Fig 4A, paired t-test, t_64_ = 7.406, P < 0.001). However, social rank influence is manifested as the differential RSTs: as social rank decreases, RST starts to decrease (Fig 4B, one-way repeated ANOVA, F(2,128) = 13.317, P < 0.001). Since RST is calculated as the difference between the start time of Δ*M_s_* and Δ*M_o_*, the decrease of RST suggests that the lower social rank facilitates faster processing of the social attribute (other’s payoff). On the contrary, better social rank prompt subjects to process private attribute (own payoff) more immediately.

**Fig 4.**
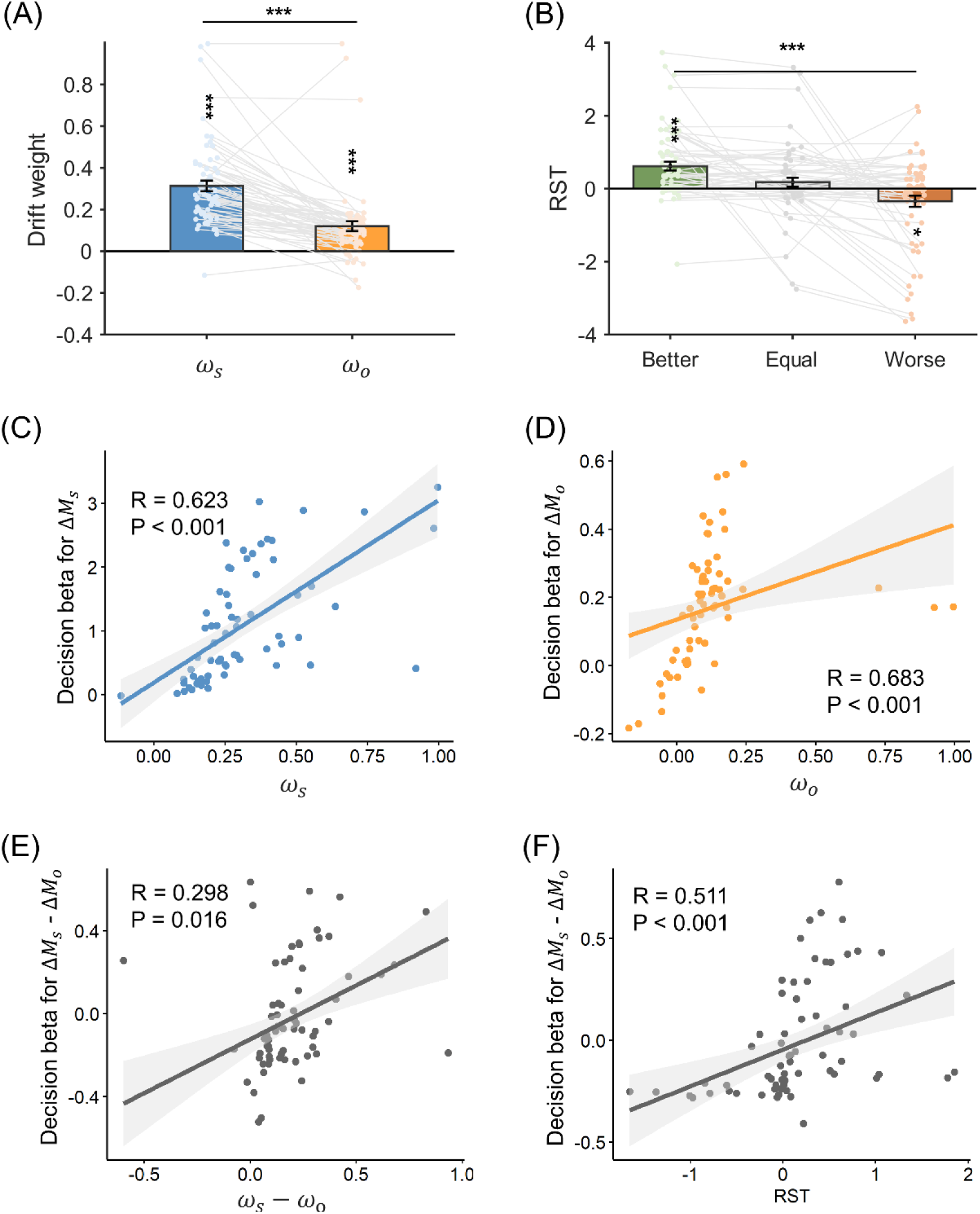
mtDDM parameters and correlations with decision betas. (A), Model estimated drift weight. (B), Model estimated RST. (C-D), Correlations between drift weights (𝜔_𝑠_ and 𝜔_𝑜_) and decision betas for Δ*M_s_* (C) and Δ*M_o_* (D) after controlling for the effect of RST. (E), Correlation between the relative drift weight (Δ*M_s_* - Δ*M_o_*) and relative decision beta (Δ*M_s_* - Δ*M_o_*), controlling for the effect of RST. (F), Correlation between RST and relative decision beta, controlling for the effect of relative drift weight. ***p < 0.001, * p < 0.05. Error bars represent s.e.m across subjects.

Interestingly, this model captures not only social-rank related behavioral differences, but also inter-subject behavioral variance across different social rank. Decision betas of Δ*M_s_* and Δ*M_o_*, the model-free estimates of the influence of Δ*M_s_* and Δ*M_o_* on choice behavior, show strong correlation with model-estimated drift weights of Δ*M_s_* (Fig 4C, robust correlation, R = 0.623, P < 0.001) and Δ*M_o_* (Fig 4D, R = 0.683, P < 0.001) respectively across social ranks as well as within each rank (S5A-F Fig). Similar correlations are also observed when the relative decision betas between Δ*M_s_* and Δ*M_o_* is regressed against the relative drift weights of Δ*M_s_* and Δ*M_o_* (Fig 4E, robust correlation, R = 0.298, P = 0.016), or against RSTs (Fig 4F, R = 0.511, P < 0.001) both across social rank and within each rank condition (see S5G-I Fig for details). The above results suggest that rank-dependent RSTs are responsible for both the social-rank related behavioral variance (Fig 2) and the inter-subject behavioral difference across conditions (Fig 4F). Across subjects, those whose social attributes (Δ*M_o_*) enter earlier into the decision process show relatively higher behavioral influence by Δ*M_o_* (Fig 4F). Additionally, there is a significant interaction effect between relative (Δ*M_s_* − Δ*M_o_*) drift weights and RSTs on the relative decision betas (𝛽 = 0.734, 𝑝 = 0.008), indicating that RST also exhibits a modulation effect of the drift weights on the decision betas, amplifying the influence of drift weights on subjects’ actual choice preference.

Given RSTs’ explanatory power in explaining relative decision betas within (Fig 2B-C) and across (Fig 4F) social rank, we further hypothesize that RSTs might directly mediate the correlation between social rank and subjects’ prosocial preferences. Therefore, we ran a mediation analysis and it revealed that RSTs partially mediated the effect of social rank on the relative (Δ*M_s_* − Δ*M_o_*) decision betas (Fig 5A, c= −0.311, P < 0.001; c^′^ = −0.213, P < 0.001) after controlling for the effects of drift weights. Similarly, RSTs also partially mediated the effect of social rank on subjects’ prosocial preference measured either by selecting options with larger *M_s_* (Fig 5B, c= −0.347, P < 0.001; c^′^ = −0.169, P = 0.007) or options with larger *M_o_* (Fig 5C, c = 0.353, P < 0.001; c^′^ = 0.228, P < 0.001), again controlling for the confounds of drift weights’ effects. These results further confirm the key role of RSTs in determining subjects’ prosocial preference.

**Fig 5.**
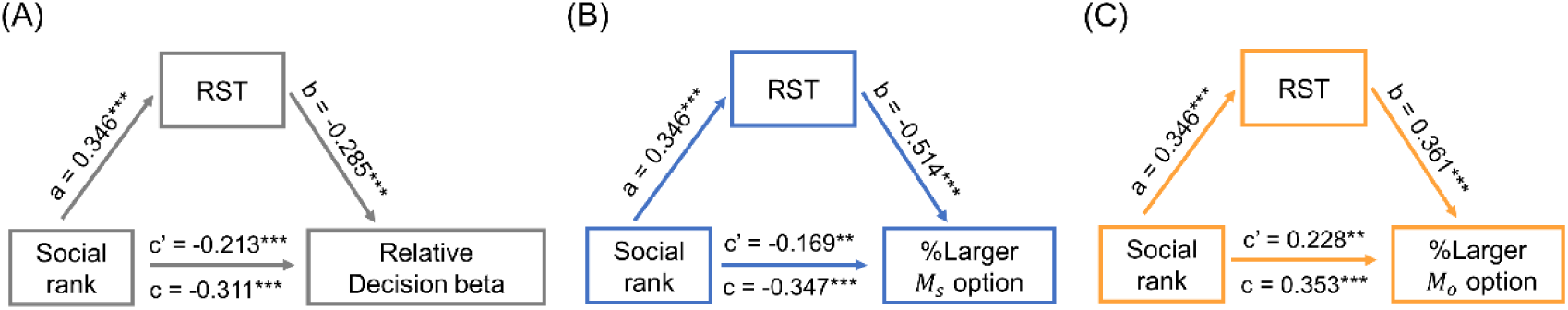
Mediation analysis. (A) RST partially mediated the effect of social rank on subjects’ relative decision beta (Δ*M_s_* - Δ*M_o_*). Similarly, RST partially mediated the effect of social rank on subjects’ preference for chosen larger *M_s_* (B) or larger *M_o_* (C) options.

### Social rank and prosociality on prosocial preference

Since social rank has a direct effect on people’s prosocial behavior and the degree of prosociality has been shown to be a reliable predictor of subjects’ prosocial behavior [47,48], we then examined whether the social rank related prosocial preference was modulated by subjects’ prosociality. To test this, we first divided subjects into prosocial (N=35) and individualistic (N=30) groups based on their social value orientation [49] (SVO) categorization criteria (see methods) and then compared the prosocial choice and RT differences of the two groups. A model-free two-way ANOVA (SVO type (2)×social rank (3)) of choice proportions revealed that the prosocial group was less likely to maximize self-payoff (option with larger *M_s_*, S6A Fig, SVO main effect: F(1,63) = 8.643, P = 0.005) and more likely to choose options with larger *M_o_* (S6B Fig, SVO main effect F(1,63) = 12.458, P = 0.001), in addition to the significant main effects of social rank reported earlier (Fig 2A). Interestingly, the interaction effects of the social rank and SVO were also significant in their choice preferences (S6A Fig, F(2, 126) = 3.691, P = 0.028; S6B Fig, F(2, 126) = 10.999, P < 0.001), suggesting that the social rank might exert different effects on subjects’ social preference contingent on their prosociality. Similar main effects of SVO and the interaction of SVO and the social rank were also observed on the decision betas of Δ*M_s_* (S6C Fig, SVO main effect: F(1,63) = 15.607, P < 0.001; interaction effect: F(2, 126) = 12.028, P < 0.001) and Δ*M_o_* (S6D Fig, SVO main effect: F(1,63) = 1.322, P = 0.255; interaction effect: F(2, 126) = 5.836, P = 0.004).

Our model-based analysis further confirmed the results above. For both SVO groups, we ran Bayesian model comparison separately and the RST model (Model 2) outperformed all the other candidate models in both groups (S2B-C Fig, prosocial group: EXP = 0.941; individualistic group: EXP = 0.999). Best fitting model parameters in the RST model showed the general pattern of the drift weight bias between attributes Δ*M_s_* and Δ*M_o_* in both groups (Fig 6A, prosocial group: t_34_ = 5.528, p < 0.001; individualistic group: t_29_ = 4.123, p < 0.001). However, the interaction effect of SVO (prosocial vs. individualistic) and attribute (Δ*M_s_* vs. Δ*M_o_*) types was also significant (F(1, 63) = 8.527, p = 0.005), indicating an even more biased attribute accumulation process in the individualistic group, which was also confirmed by the slower RTs in the prosocial group (S6E Fig, F(1,63) = 17.540, P < 0.001). Lastly, we examined how the social rank and SVO influenced subjects’ RST in a two-way ANOVA. Although the SVO type did not yield a main effect on RST (Fig 6B, F(1,63) = 0.046, P = 0.832), the SVO and social rank interaction was significant (F(2,128) = 3.669, P = 0.028), suggesting that the individualistic group’s RST was less affected by the social rank. Indeed, closer examination of each SVO group showed that the social rank effect was significant in the prosocial group (F(2, 68) = 11.153, P < 0.001) but only reached marginal significance in the individualistic group (F(2, 58) = 2.938, P = 0.061). Taken together, these results demonstrate that the RST model not only captures the behavioral patterns of individual differences across subjects (Fig 2), its validity is also reflected in both the prosocial and individualistic groups. Similarly, our simulation results confirm that the RST model faithfully predicts the behavioral choice pattern and response time in both groups (S6 Fig, red error bars).

**Fig 6.**
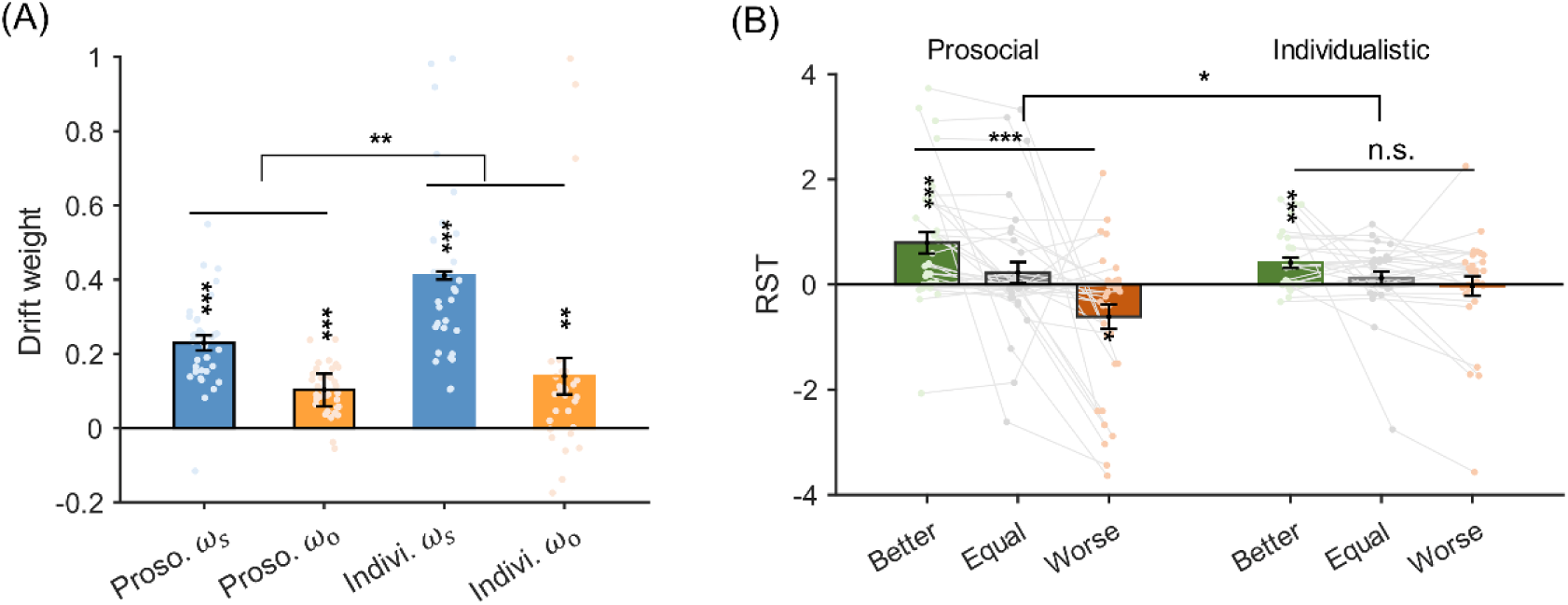
Model estimated individual differences. (A) Significant interaction indicates the relative drift weights (𝜔_𝑠_ − 𝜔_𝑜_) of the individualistic subjects are higher than that of the prosocial subjects. Asterisks across prosocial and individualistic groups denote significant interaction effect of two-way ANOVA (attribute type × SVO group). (B) Prosocial group’s RST is more sensitive to the change of social rank than that of the individualistic subjects. Asterisks across better, equal and worse social rank denote significant main effect of one-way ANOVA. *P<0.05, **P<0.01, and ***P<0.001. Error bars represent s.e.m across subjects.

### Confirmation of model validity in the replication study

To further verify the validity of our results and the reliability of the RST model in accounting for the social rank effects, we replicated the study with a lager sample size (105 subjects in total, 52 prosocial subjects and 53 individualistic subjects, see methods). In this study, we replicated all the major findings from the original study (Fig 7, S7-13 Fig, S1-3 Table): social rank modulates subjects’ prosocial preference as well as their reaction times in both prosocial and individualistic groups (S7 Fig) and the RST model performed best among the candidate models (S8 Fig). Again, in the replication study, we found significantly larger drift weight of the Δ*M_s_* than the Δ*M_o_* attribute (Fig 7A, t_104_ = 12.694, P < 0.001); and RST decreased significantly as the social ranks worsened (Fig 7B, F(2,208) = 22.426, P < 0.001).

**Fig 7.**
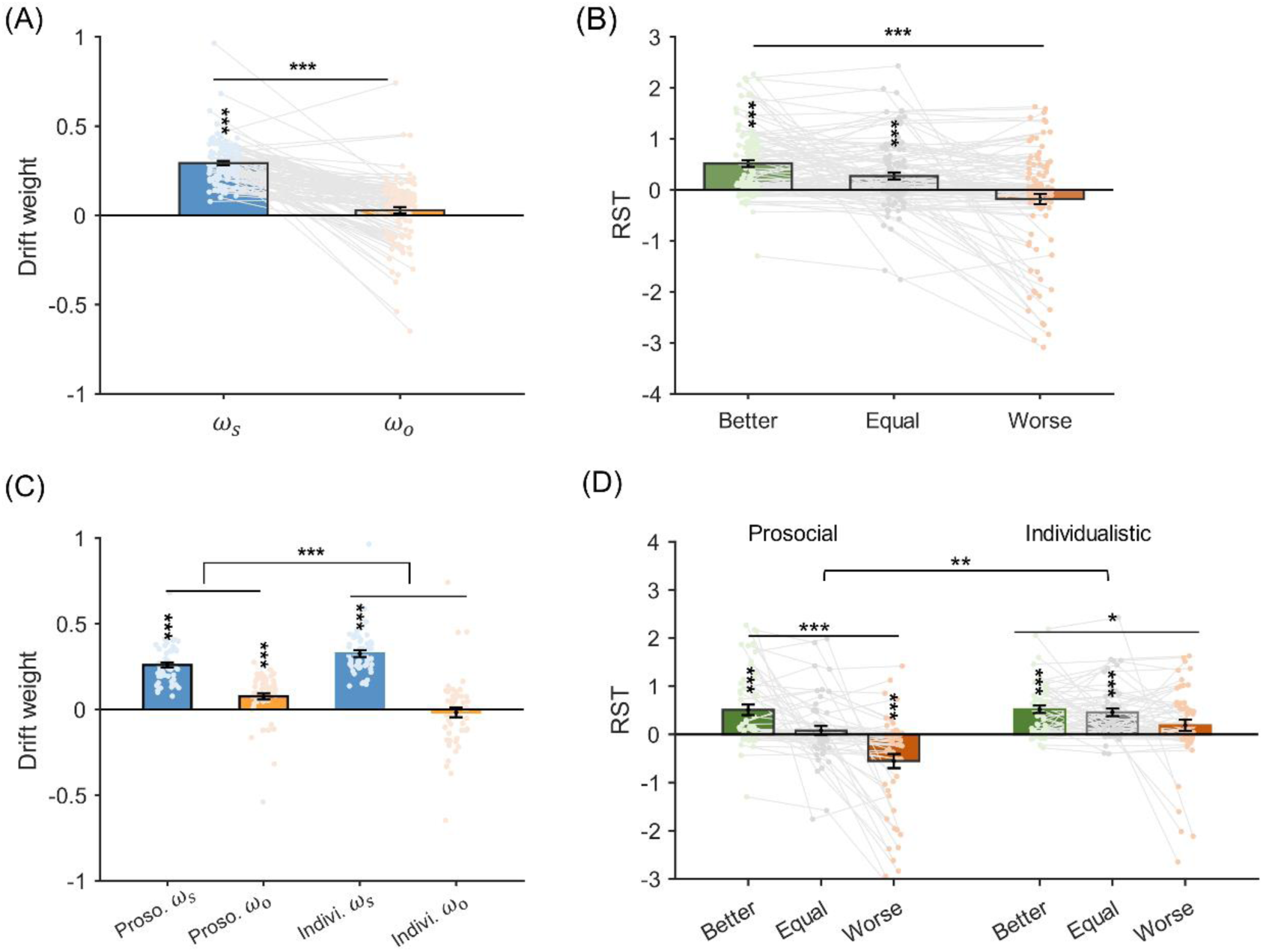
Modeling results from the replication study. (A) Model estimated drift weight. (B) Model estimated RST. (C) Prosocial and individualistic groups differ in their drift weights. Significant interaction indicates the relative drift weights (𝜔_𝑠_ − 𝜔_𝑜_) of the individualistic subjects are higher than that of the prosocial subjects. Asterisks across prosocial and individualistic groups denote significant interaction effect of the two-way ANOVA (attribute type × SVO group). (D) Prosocial group’s RST is more sensitive to the change of social rank than that of the individualistic subjects. Asterisks across better, equal and worse social rank denote significant main effect of one-way ANOVA. *P<0.05, **P<0.01, and ***P<0.001. Error bars represent s.e.m across subjects.

Furthermore, the two-way ANOVA also revealed a more biased processing of self-payoff (Δ*M_s_*) than the other-payoff (Δ*M_o_*) attribute in the individualistic group (Fig 7C, interaction effect: F(1,103) = 16.733, P < 0.001). Most importantly, the two-way ANOVA of the RST model demonstrated a significant interaction effect of social rank and SVO type (Fig 7D, F(2,206) = 6.308, P = 0.002), confirming similar attribute timing patterns in the replication study groups: compared to the individualistic group, the prosocial group’s RSTs were more sensitive to the change of social rank (Fig 7D).

### Correlation between empathy and decision process

It is generally believed that empathy signals the natural tendency to perceive others’ emotional states and motivates prosocial behavior [49–54]. We measured subjects’ empathy traits using the influential Interpersonal Reactivity Index (IRI) [55, 56], and there was a significant IRI difference between the prosocial and individualistic groups (two-sample t-test, t_157_ = 2.112, P = 0.036 across two datasets in study 1 & 2. Furthermore, across two studies, the IRI was significantly and positively correlated with subjects’ drift weight of Δ*M_o_* (𝜔_𝑜_, Fig 8A, robust correlation, R = 0.205, P = 0.010), but not the drift weight of Δ*M_s_* (𝜔_𝑠_, Fig 8B, robust correlation, R = −0.037, P = 0.642). These results confirm previous findings that empathy carries the capacity to automatically sense and incorporate others’ affective states and therefore is linked to the motivation of prosocial preference [57–59]. In addition, we also found that the IRI was negatively correlated with the RST’s rank sensitivity, defined as the linear slope of RST across better, equal and worse social ranks (Fig 8C, R = −0.164, P = 0.039), suggesting that participants with higher empathy levels adjusted their RSTs more adaptively according to the change of social ranks. Together, these results resonate well with the motivational account of empathy and highlight both the automatic and the adaptive nature of empathy [57,60].

**Fig 8.**
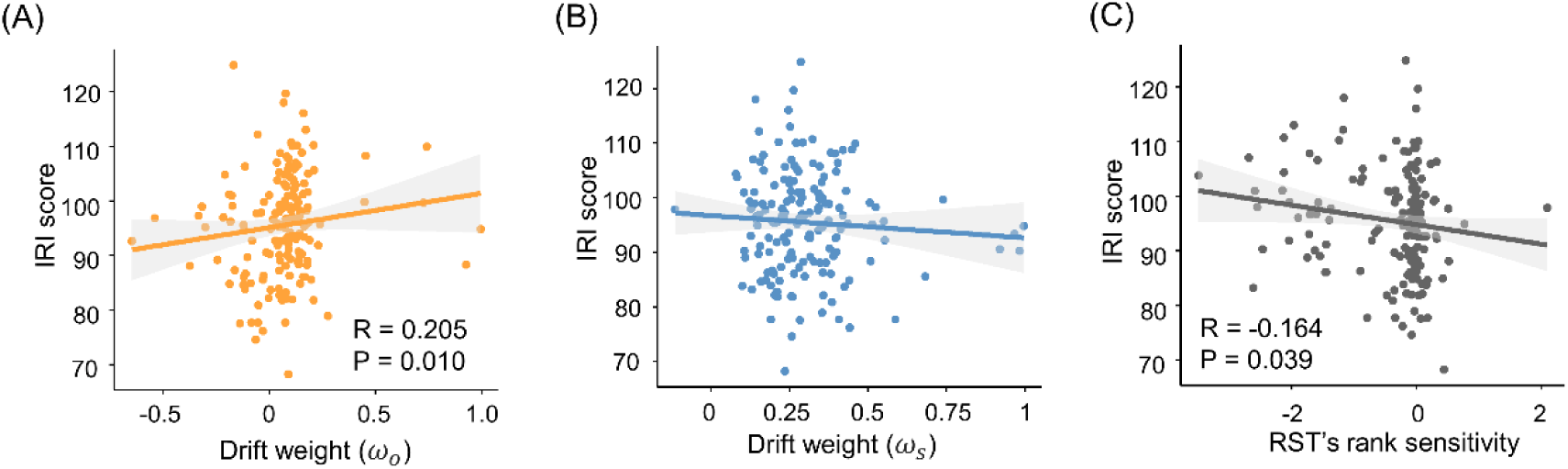
Correlation between IRI score and model parameters. (A) IRI score is significantly correlated with drift weights for others’ payoffs (𝜔_𝑜_). (B) However, IRI score showed no correlation with drift weights of their own payoffs (𝜔_𝑠_). (C) Significant correlation between IRI score and subjects’ RST rank sensitivity (individual linear slope of RST across social ranks).

## Discussion

Do social rank influence people’s generosity? Our results suggest they do, but in a more nuanced manner. With the combination of social rank manipulation and computational modeling, our results provide empirical support to delineate two possible venues via which social rank might influence prosocial preference: when and how strongly the private (Δ*M_s_*) and social (Δ*M_o_*) attributes affect decisions. We have shown that social rank modulation was specifically associated with the relative start time (RST) of the decision attributes entering into the decision process: as social rank worsened, the social attribute was contemplated first before the private payoff attribute was taken into consideration. Such asynchrony can ostensibly manifest itself as higher decision betas (higher decision regression coefficients) to the social attributes and results in the more prosocial preference in the lower social rank condition. However, with the aid of the formal model of decision processes, we show that social rank is closely related to the attribute timing. Interestingly, attribute drift weights are spared by the social rank change. Furthermore, the social rank effects are partially mediated by the RST, indicating that attribute timing plays an important role in explaining subjects’ altruistic behavior (Fig 5).

Previous literature on social rank and morality have shown that higher social class is a good predictor for unethical behaviors including lying in negotiations, cheating and law breaking [61,62]. Subsequent studies further demonstrate that context specific social rank such as winning a competition also prompts subjects to behave more dishonestly [63], indicating that both stable and dynamic social rank can predict moral choices. Conversely, people are averse to the transgression of the established social ranks via social welfare distribution. For example, it has been shown that despite social inequality aversion, people accept more unequal offers in a money allocation choice game to avoid reversing the ranks of social agents [36]. What remains unaddressed is how the prosocial preference, usually considered as a stable trait [49], can be modulated by the change of experimentally induced social rank [64]. One possibility is offered by the query theory which suggests that valuation is constructed by posing a series of queries differing in content and sequence idiosyncratically [37,38,65]. Our asynchronous attribute accumulation approach can be viewed as the mechanistic implementation of the query theory: the malleability of prosocial preference in response to the change of social rank is reflected in the order of decision attributes taken into consideration during the decision process. Alternatively, it is also conceivable that attribute weights are subject to the specific decision context. For instance, recent studies in value-based decisions have shown that explicitly instructing subjects to pay attention to the tastiness or healthiness of the food options only changes the attribute weighting strength, but not attribute timing [23]. Similar results were achieved via neural interventions such as transcranial direct current (tDCS) stimulation over the dorsolateral prefrontal cortex (dlPFC) in the food choice task [22]. In the social decision domain, it has been shown that the dynamic attribute weighting DDM guided by the eye-gaze informed attention better explains participants’ social preference and response time under time pressure [12]. These studies suggest that both the query content and sequence can be adaptively modulated depending on the specific decision context. Our study further develops this idea and shows that the social rank has a selective effect when biasing subjects’ altruistic choices: lower social rank specifically leads to sooner consideration of social attribute.

Our results shed light on an important aspect of social decision-making. By leveraging both RT and choice behavioral data, we utilize a DDM with asynchronous attribute processing and provide a mechanistic account for the impact of social rank on altruistic choices that cannot be fully accounted for by static decision models alone. Importantly, such findings were further corroborated in an independent and larger dataset, validating the robustness of our approach. Previous studies concerning social rank manipulation have documented flexible social preference in different contexts [66,67]. The selective effect of social rank on attribute timing may stem from conflicts between situational or context-specific social norm and subject’s internal social preference [68,69]. For instance, the inclination towards prosocial preference in reward allocation might not be suitable in the high social rank context, where social norm dictates that one should receive a larger portion of the pie compared to their co-players. Consequently, it is plausible that the attribute timing adjustment serves as a proactive strategy to shape behavior, potentially aided by the flexible allocation of attention [70,71], while leaving subjects’ intrinsic social preferences unchanged.

The social value orientation (SVO) theory distinguishes between prosocial and individualistic participants based on their attitudes towards the well-being of others [72], yet it remains agnostic about the malleability of prosociality in response to the changes in decision contexts. The empathy account of prosocial preference underscores people’s inherent inclination to perceive and align with the emotional states of others, motivating concerns for others’ welfare [50–52,54]. Our findings are consistent with previous studies by showing that the prosocials are indeed inherently more generous than the individualists (Figs 6A and 7C). More importantly, we extend the SVO theory by revealing that prosocial individuals also exhibit greater adaptability in response to changes in social rank, in contrast to the individualists (Figs 6B and 7D). It suggests that in addition to being more “altruistic”, the prosocials are also more sensitive to the decision context-specific social norm than the individualists [68,69]. Intriguingly, these results dovetail nicely with the motivated account of empathy, which posits that empathy as a multi-facet construct encompasses different subcomponents [55–57,62]. The automatic facet of empathy, which comprises experience sharing, mentalizing and mind perception, may be the root of the “baseline” level of prosociality. Indeed, we find that the IRI score, an influential measure of empathy, is positively correlated with subjects’ drift weight toward others’ payoff (Δ*M_o_*) but no correlation exists between IRI score and the drift weights of subjects’ own payoff (Δ*M_s_*, Figs 8A&B). On the other hand, empathy varies with different situations. In our study, the IRI score also correlates with how sensitive RST adapts to different social rank (Fig 8C), underscoring the context dependency aspect in empathy [57]. Hence, the classification of subjects based on SVO and their attribute timing sensitivity towards social rank can be unified under the framework of motivated empathy, implying a novel dimension through which social rank, prosociality and empathy can be connected [61,62].

There are also limitations in our approach that warrant further investigation. First, we adopted an RST DDM where attribute onset was relative to each other instead of an independent mtDDM that allows separate time onsets for different attributes [22, 23]. In future research aimed at modeling decisions involving multiple attributes, it will be imperative to expand the current model to the independent mtDDM. Furthermore, there has been clear evidence that social preference is a dynamic construct significantly influenced by the moment to moment attention deployment during the decision process [12]. Indeed, results from a recent eye-tracking study suggest that the SVO difference is associated with the degree of attention directed towards the other’s payoffs in a money allocation task, highlighting the intertwined nature of social preference and attention allocation [18]. Future studies could benefit from combining the eye-gaze informed attention mechanism with the assumption of asynchronous attribute timing in explaining social behavior. Lastly, while the DDM has garnered substantial support from neural evidence [73–76], the brain mechanism of attribute timing remains elusive. Recently, it has been reported that the cathodal tDCS stimulation to the left dlPFC only altered subjects’ weighting strength of the tastiness attribute but not the attribute timing. Future high temporal resolution electroencephalogram (EEG) and magnetoencephalogram (MEG) studies, in combination with representation encoding analysis might be required to study such a dynamic process where the change of RST is in the range of sub-seconds [48].

In conclusion, our results demonstrate that the adaptation of social preference to specific social context can be accounted for by the asynchronous attribute timing in the decision process. By extending the mtDDM from value-based decision paradigms to the social decision domain, we show that the adjustment of attribute start time (or RST) may be a general mechanism for inducing contextually congruent behaviors without altering underlying preferences. Our findings also highlight the intrinsic differences among individuals with different prosociality and empathetic concern: not only they differ in their social decision attribute weights, but also differ in the RST adaptation sensitivities in response to changes in social rank. These findings may prove useful in devising public policies and interventions that encourage faster rather than stronger social attribute processing in promoting prosocial behaviors.

## Methods

### Subjects

In the original study (study1), we recruited 65 healthy college adult students (42 females, age = 22.357 ± 2.526 years old; 23 males, age 22.826 ± 2.480). In the replication study (study 2) we recruited 114 healthy college adult students (70 females, age = 20.914 ± 2.418 years old; 44 males, age 21.273 ± 2.815) and asked them to rate how much they believe to play with a real partner (Likert scale 0-5). 9 subjects were excluded due to their disbelief of the experimental settings (Likert rating of 0), hence data of the remaining 105 subjects were used in further analyses. The experiment was conducted in accordance with the protocol approved by the Ethics Committee of School of Psychological and Cognitive Sciences, Peking University. Informed written consent was obtained from each subject before each experiment.

### Task

We used a two-stage decision task combining a dot number estimation game and a dictator game (DG, subjects as the money allocators) (Fig1 A). In study 1, upon arrival, subjects were instructed to play with an anonymous co-player (confederate) who finished the same task previously, and they would have the chance to pair with future subjects and receive payoffs allocated by the future subjects. In each trial subjects were first asked to choose which of the two options on the screen contained more grey dots (the dot number estimation game) within 3 seconds, after which they received performance feedback of both players for 2 seconds. Then in the dictator game, subjects had to choose between two payoff allocation options for themselves (*M_s_*) and the co-players (*M_o_*) with no response time limit imposed. After the task, subjects were informed that all the payoff in the chosen options would be summed up and converted to real money for them and the co-players to keep, and if they were selected as co-players for the future subjects, they would receive an extra amount of payoff as the co-players.

To construct a more realistic social decision environment, we recruited cohorts of 6-12 subjects each in the replication study. They were instructed that in each trial they would be randomly and anonymously paired with other subjects in the cohort. Each trial started with the dot number estimation game and after the performance feedback, subjects were asked to finish 2-4 DG choice games with the maximum reaction time of 6s (1.026% of trials exceeded RT window of 6s in study 1) for each DG game. All the other settings were the same as study 1.

In study1, the whole task consisted 5 sessions (60 trials each), and subjects on average took 50 minutes to finish the task. In the replication study, the whole task was separated into 4 sessions (25 dot number estimations each), and it took about 35 minutes to finish the task. Subjects received payoff ranging from ¥55-75 (RMB) in both studies according to their choices (¥50/hour show up fee + choice related payoffs).

For the dot number estimation game, the total dots associated with each option were randomly drawn from a uniform distribution of [32,34] with the constraint that dots numbers of two options are not equal. Our behavioral results confirmed that subjects’ performance was at chance level (accuracy = 0.5) with reaction time window of 3s (study 1: mean accuracy rate = 0.506, one sample t-test, t_64_ = 1.698, P = 0.047; replication study: mean accuracy rate = 0.504, t_104_ = 1.109, P = 0.155;). Unbeknownst to the subjects, we manipulated the performance feedback such that the number of trials for each feedback type was 60 (both players were correct, BC), 60 (both were wrong, BW), 90 (subject correct and the partner wrong) and 90 (the partner correct & subject wrong), respectively. Since there was no difference in the prosocial preference of the BC and BW trials (S1 Fig), we further collapsed trials of BC and BW types (equal social rank group) to organize subjects’ performance relative to their partners and thus resulting in better (90 trials), equal (120 trials) and worse (90 trials) social rank conditions (Fig 1B).

For the binary dictator game, in each option the monetary payoffs for the subject (*M_s_*) and the co-player (*M_o_*) were randomly drawn from 1 to 20 points with the following constraints: (1) The correlations coefficients among 𝑀_𝑠1_, 𝑀_𝑜1_(option 1), 𝑀_𝑠2_, 𝑀_𝑜2_ (option 2) were between -0.1 and 0.1. (2) The correlation coefficients between Δ*M_s_* (𝑀_𝑠1_*-*𝑀_𝑠2_) and Δ*M_o_* (𝑀_𝑜1_*-*𝑀_𝑜2_) were between -0.1 and 0.1 to avoid collinearity among variables.

### Behavioral analysis

To investigate the influence of social rank on subjects’ choices, we calculated proportion of trials where subjects chose either the larger *M_s_*, or the larger *M_o_* options respectively (Fig 2A, S6A&B Fig, and S7A-C Fig). We also applied a linear mixed model (LMM) to examine whether social rank was associated with how Δ*M_s_* and Δ*M_o_* affected subjects’ choices:

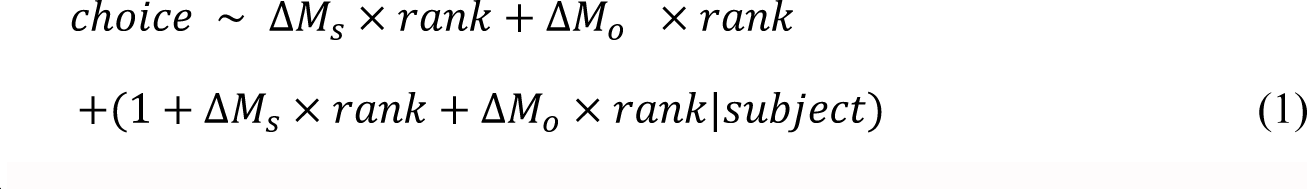

Where 𝑠𝑡𝑎𝑡𝑢𝑠 is a factor coding for the social rank (better, equal or worse), and the choice was coded as 1 if subjects chose the larger *M_s_* options and 0 otherwise. The results are presented in Fig 2B&C, S6C&D Fig, and S7D-F Fig.

### Computational Modeling

To fit subjects’ choice and RT, we employed a multi-attribute and time-varying drift diffusion model (mtDDM). This model assumes that different attributes may enter the evidence accumulation process at different time onsets. Therefore, the decision-making process (choice and RT) can be influenced by both the attribute drift weights and their corresponding starting time. The decision evidence accumulates in discrete time (𝑑𝑡), and the evidence accumulation stops when the decision variable reaches the decision boundary or threshold.

At each time point 𝑡, the drift rate (𝜈_𝑡_) is governed by the attributes strength (𝜔_𝑠_ for Δ*M_s_* and 𝜔_𝑜_ for Δ*M_o_*, respectively) as well as the drift latency parameters *𝜏^𝑡^_s_* and *𝜏^𝑡^_o_*, which are associated with the relative start time (RST) between attributes.

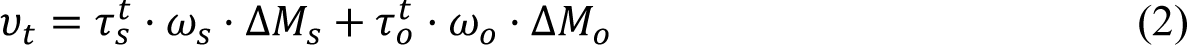

Specifically, if 𝑅𝑆𝑇 > 0, Δ*M_s_* enters the decision process first; else if 𝑅𝑆𝑇 < 0, Δ*M_o_* enters first. Finally, if 𝑅𝑆𝑇 = 0, the two attributes enter the decision process simultaneously:

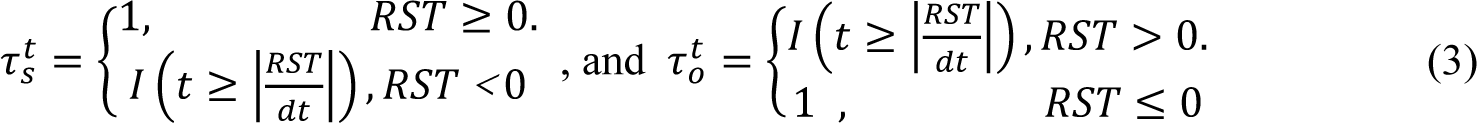

Where 𝐼(·) is an indicator function.

The change of social rank may influence the decision process either by modulating the attribute strength (𝜔_𝑠_ and 𝜔_𝑜_) or by adjusting the relative start time (RST) between attributes. We therefore constructed a series of candidate models and tested their performance against our behavioral data (S1 Tables).

**Model 1.** Social rank may have specific effects on the attribute strength and both attributes enter the decision process at the same time (𝑅𝑆𝑇 = 0):

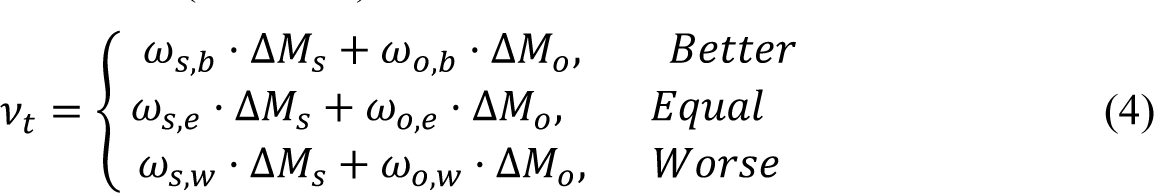

Where the 𝜔_𝑠,𝑏_, 𝜔_𝑜,𝑏_, 𝜔_𝑠,𝑒_, 𝜔_𝑜,𝑒_ and 𝜔_𝑠,𝑤_, 𝜔_𝑜,𝑤_ are the attribute strength of Δ*M_s_* and Δ*M_o_* in the better, equal and worse social rank, respectively (Eq. 1).

**Model 2.** The social rank may only modulate the RST but not the attribute strength.

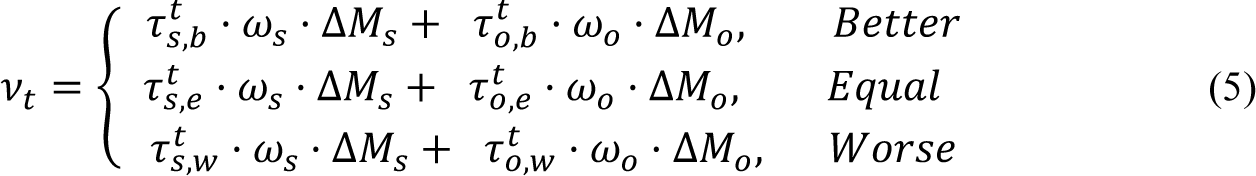

Where *𝜏^𝑡^_s,b_*, *𝜏^𝑡^_o,b_*, *𝜏^𝑡^_s,e_*, *𝜏^𝑡^_s,w_*, *𝜏^𝑡^_o,w_*, 𝜏^𝑡^ are determined by RSTs (𝑅_*b*_𝑆𝑇_*e*_, 𝑅𝑆𝑇_*w*_ 𝑎𝑛𝑑 𝑅𝑆𝑇)

for different social ranks (better, equal or worse), respectively (Eq. 2).

**Model 3.** We further assume that the RST is constant across social ranks but the attribute strength is not (Eq. 3):

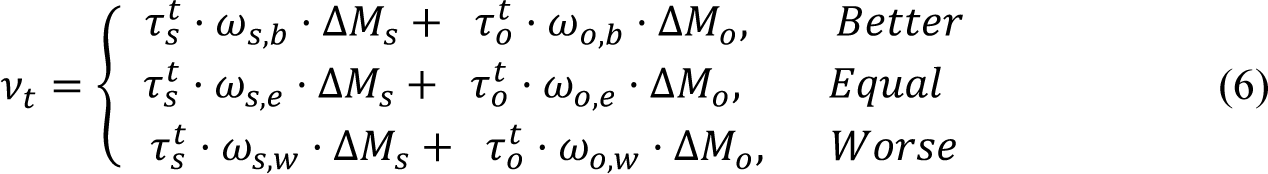

**Model 4.** We assume that social rank modulates both the attribute strength and the relative start time of the attributes (Eq. 4).

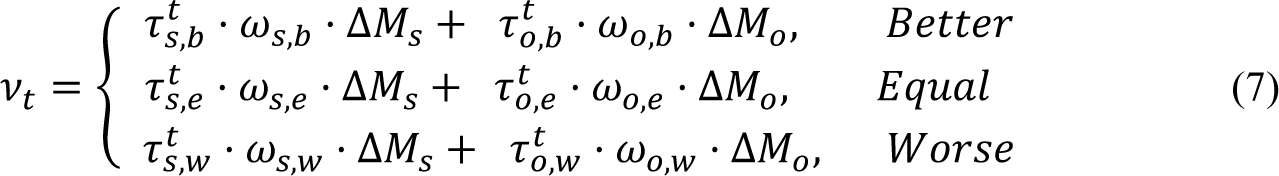

Where the drift latency and attribute strength parameters are the same as previously defined. We also include the following parameters in the model: (1) Decision boundary (Threshold):

the evidence threshold for the decision, which is set to be symmetric around zero. (2) nDT: the non-decision time, which accounts for the extra amount of time required for subsequent motor action not related to evidence accumulation. (3) Bias: the starting point bias for choosing larger *M_s_* option in the evidence accumulation process.

With the drift rate (𝜈_𝑡_) specified, the evidence-updating equation is as follows (Fig 3A & B):

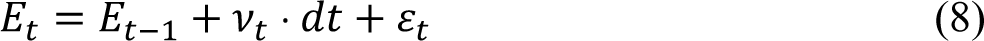

Where 𝜀_𝑡_ represents an i.i.d. Gaussian noise term from 𝑁(0, 0.1). Evidence accumulation proceeds as follows. It begins with the initial bias (𝐸_0_ = 𝐵𝑖𝑎𝑠). It is then updated in discrete time steps of 𝑑𝑡 (0.01𝑠), until | 𝐸_𝑡_ | reaches the decision boundary and the response time (RT) is computed as 𝑡 · 𝑑𝑡 + 𝑛𝐷𝑇.

For each candidate model and each subject, we estimated the best-fitting parameters using the differential evolution algorithm (an R Package developed by Mullen and colleagues [44]) with 150 iterations. In each iteration, we simulated 3,000 choices and RTs for each trial. Trials with actual RT longer than 10s and beyond three standard deviations were excluded from model-fitting, and this exclusion criteria resulted in a total of 52 trials discarded for all the subjects (less than 0.3% of all trials in study1, and no trial was discarded in the replication study). The likelihood of the observed choices and RTs were generated according to the distribution generated by the 3000 simulated choices. For each subsequent iteration, the parameter populations evolved towards maximizing the likelihood of the observed data (subjects’ choices and RTs) by the procedures described in Mullen, et al^44^. We repeated above process in each model 100 times to get the best parameter estimations.

Bayesian model selection was used for model comparison [46] (S2-8 Fig) and the candidate model performance was summarized in S2 Table (study 1) and S3 Table (replication study).

### mtDDM description

We simulated model predicted decision betas of Δ*M_s_* and Δ*M_o_* as functions of RT. We simulated each trial 100 times for all subjects’ option sets with the mean parameters of Model 2. To more intuitively show the effect of attribute weight and timing on choice, the initial bias was set to 0. Then, we ran logistic regressions (𝐶ℎ𝑜𝑖𝑐𝑒 ∼ Δ*M_s_* + Δ*M_o_*) in 100ms moving time windows in steps of 10ms. This result is shown in Fig 3C-E.

### Model parameter recovery

We ran parameter recovery for the best model by simulating choices and RTs based on the estimated parameters for each subject and each trial. Then, we separately fitted the simulated behaviors using the best model and recovered the model parameters for each subject. We then calculated the correlation between the recovered parameters and true parameters originally estimated from fitting subjects’ actual behavior. This result is shown in S3 and S10 Fig.

### Model cross-validation

To conduct the cross-validation analysis for the RST model, we randomly separated subjects’ data into two parts for each social rank condition in each subject. Next, we used the parameters estimated by half of the data to simulate subjects’ choices and RTs for the option sets of the other half data 100 times.

We then computed the simulated choice accuracy against subjects’ actual choices and the RT correlation with subjects’ true RTs in the other half data. The results are shown separately for better, equal, and worse conditions in S4 and S9 Fig.

Next, we tested how the best model could capture subjects’ behavior patterns. We averaged the simulated choices and RTs for the 100 stimulations of each trial and compared the model predicted choices in each social rank condition (Fig 2A, S6A&B Fig, and S7A-C Fig, red error bars). We also retrieved the decision regression coefficients Δ*M_s_* and Δ*M_o_* and plotted both the simulated and actual regression coefficients together for comparison (Fig 2B&C, S6C&D Fig, and S7D&F Fig, red error bars). The detailed description of this regression analysis is in the regression analysis section (see equation (1)). Finally, we plotted the simulated RTs across social rank (Fig 2D, S6E Fig, and S7G&H Fig, red error bars) along with the actual RTs.

### Questionnaires

To investigate individual differences in attribute weight and timing, we evaluated subjects’ social preference and empathy trait with the social value orientation [49] (SVO) and interpersonal reactivity index [55, 56] (IRI) questionnaires. SVO was widely used in measuring ones’ social preference. Based on SVO scores, we classified subjects into prosocial and individualistic groups (Fig 6, Fig 7C-D, S6&7 Fig). IRI is a widely used empathy measuring questionnaire that comprises 4 sections of the empathy construct, we checked how IRI related with the model results (Fig 8).

### Software and statistics

The task was presented with Psychtoolbox (3.0.17) in Matlab (2018b) on a computer screen. Computational modeling of mtDDMs was conducted using DEoptim (v2.2-6) package in R.4.1.1. Mixed effect regressions were conducted with fitglme (Matlab function). Robust correlation was calculated with Matlab (2018b) toolbox BendCorr. Mediation analysis was conducted in Matlab (2018b).

## Author contributions

J.L. and Y.N. conceived the idea and designed the experiment. Y.N conducted the experiment, performed data analysis. J.L. and Y.N. wrote the paper. All authors discussed the results and implications, and commented on the manuscript at all stages.

---

**S1 Fig.**
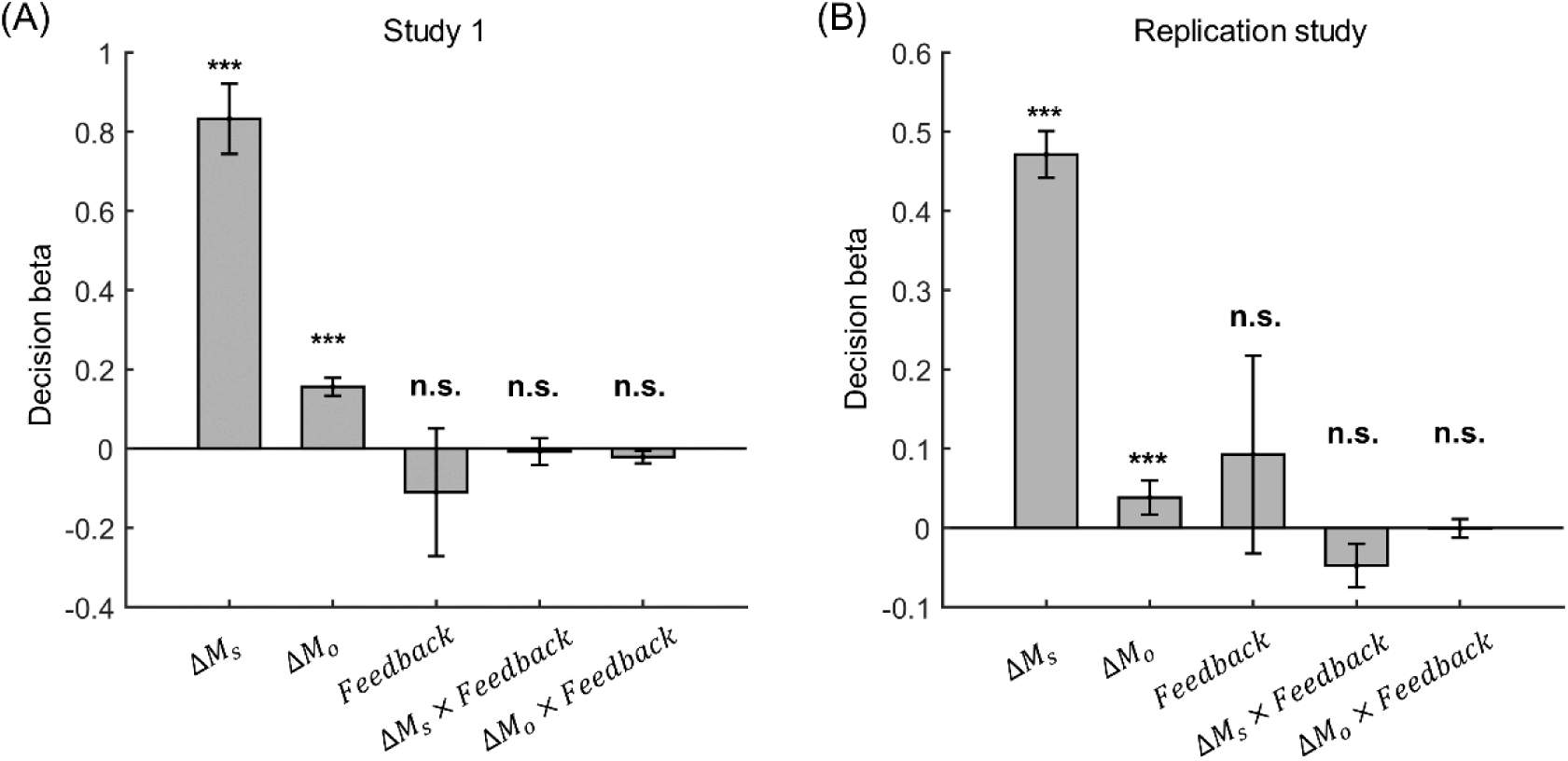
Decision regression coefficients of different attribute in the equal rank condition. Mixed-effect regression analysis results for equal social rank conditions where feedbacks from both players were correct or incorrect. The feedback was a factor coding whether both players were right or wrong in the dot estimation task. The main effects of feedback, together with the interaction effects between feedback and Δ*M_s_*/Δ*M_o_* were not significant in both studies (study 1 and replication study).

**S2 Fig.**
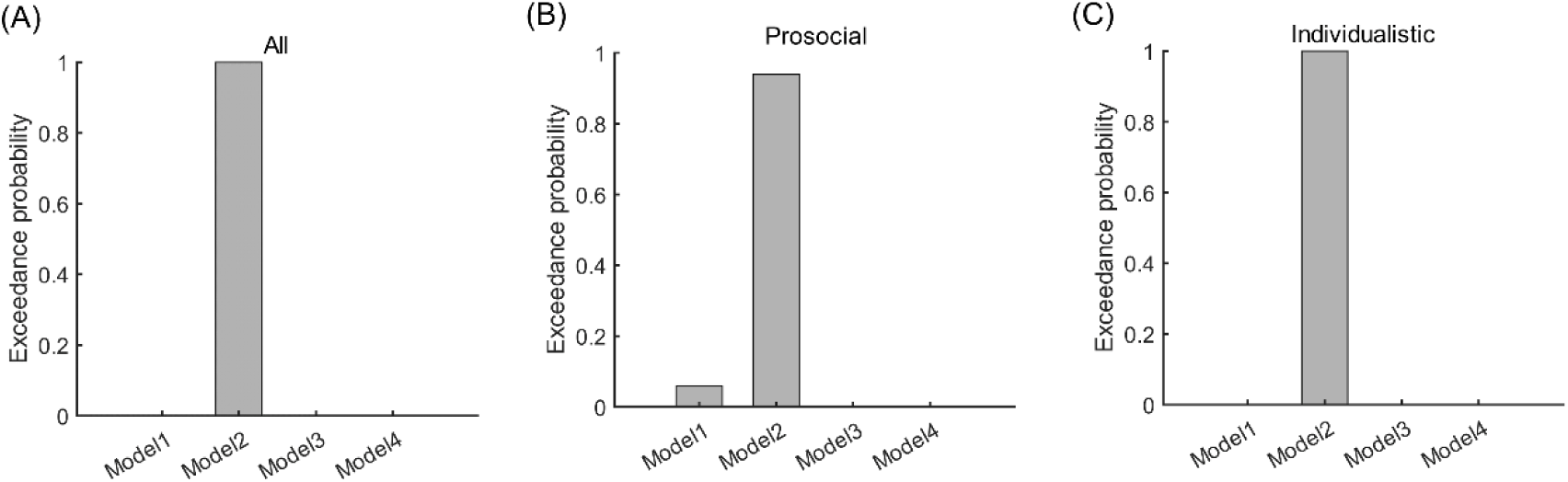
Bayesian model selection results in study 1. Across all the subjects (A) or within the prosocial group (B) or individualistic group (C), model 2 (RST model) outperformed other models with the highest exceedance probability.

**S3 Fig.**
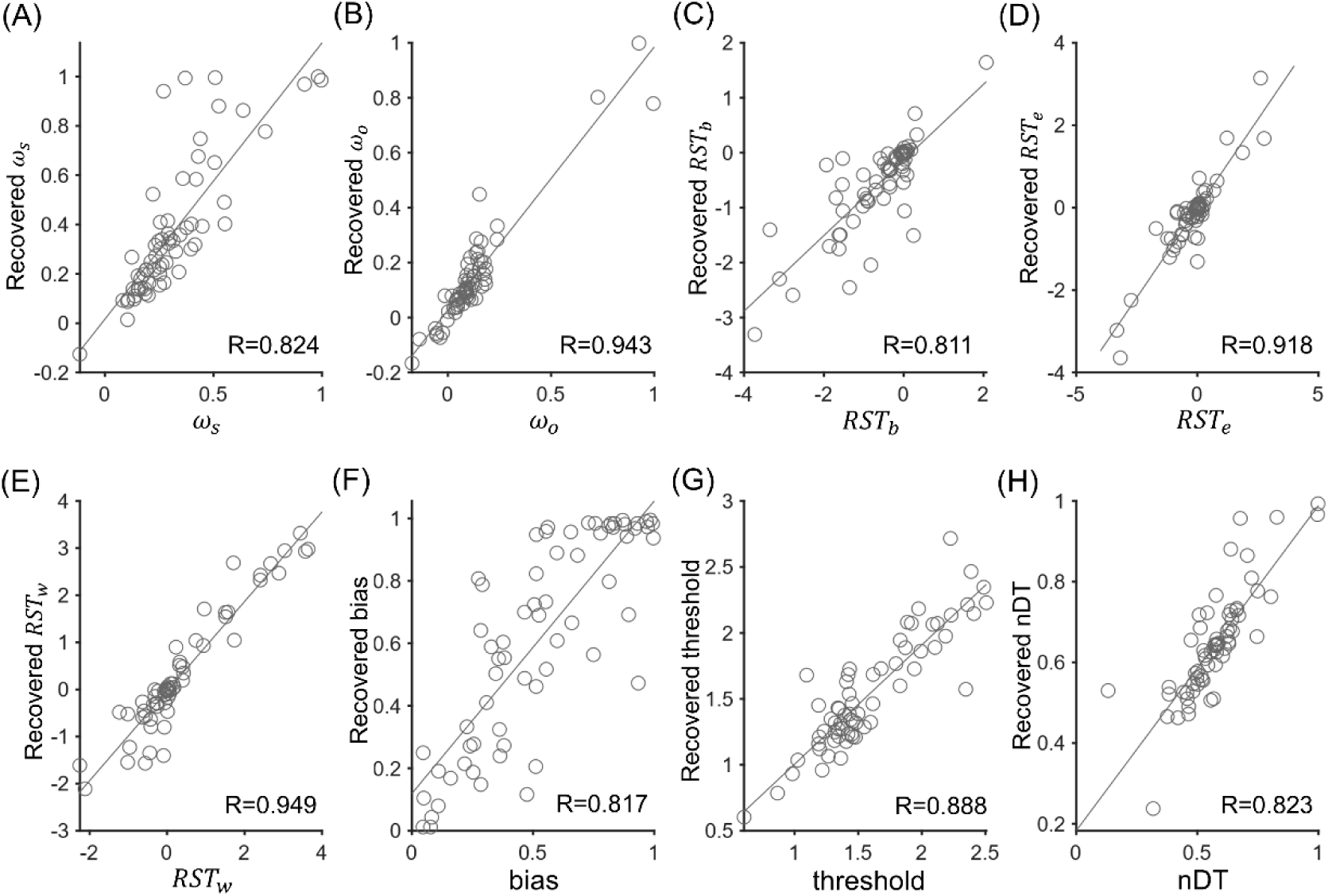
| Parameter recovery analysis for the winning RST model in study 1. Correlations between actual parameter and the model recovered parameters.

**S4 Fig.**
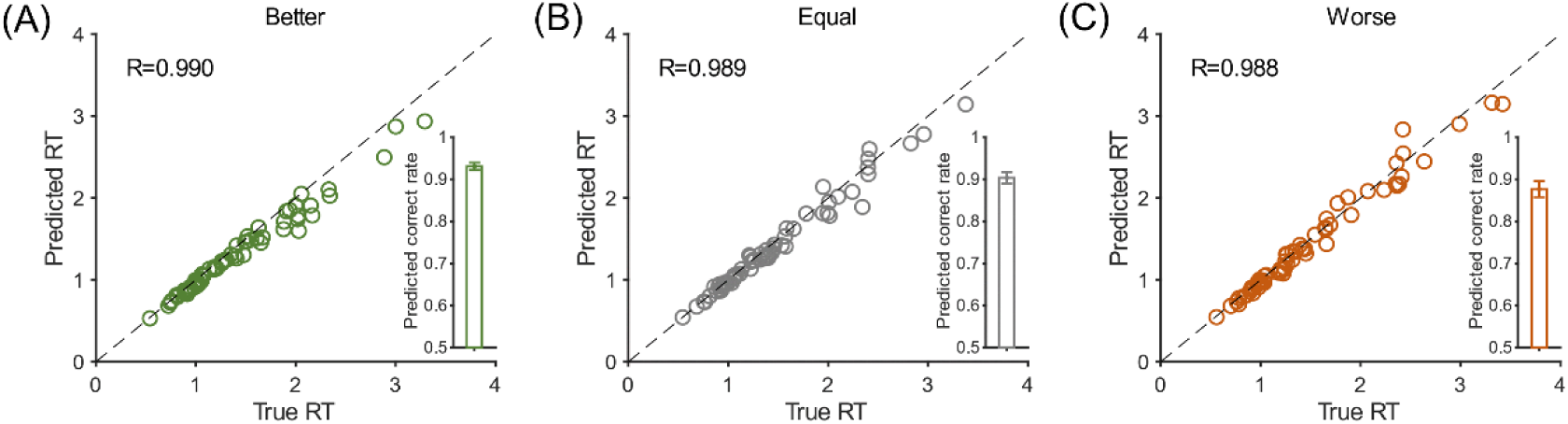
Cross-validation results in study 1. The correlation between predicted and participants’ actual RTs (empty circle) and choices (bar) across subjects in the better, equal and worse conditions.

**S5 Fig.**
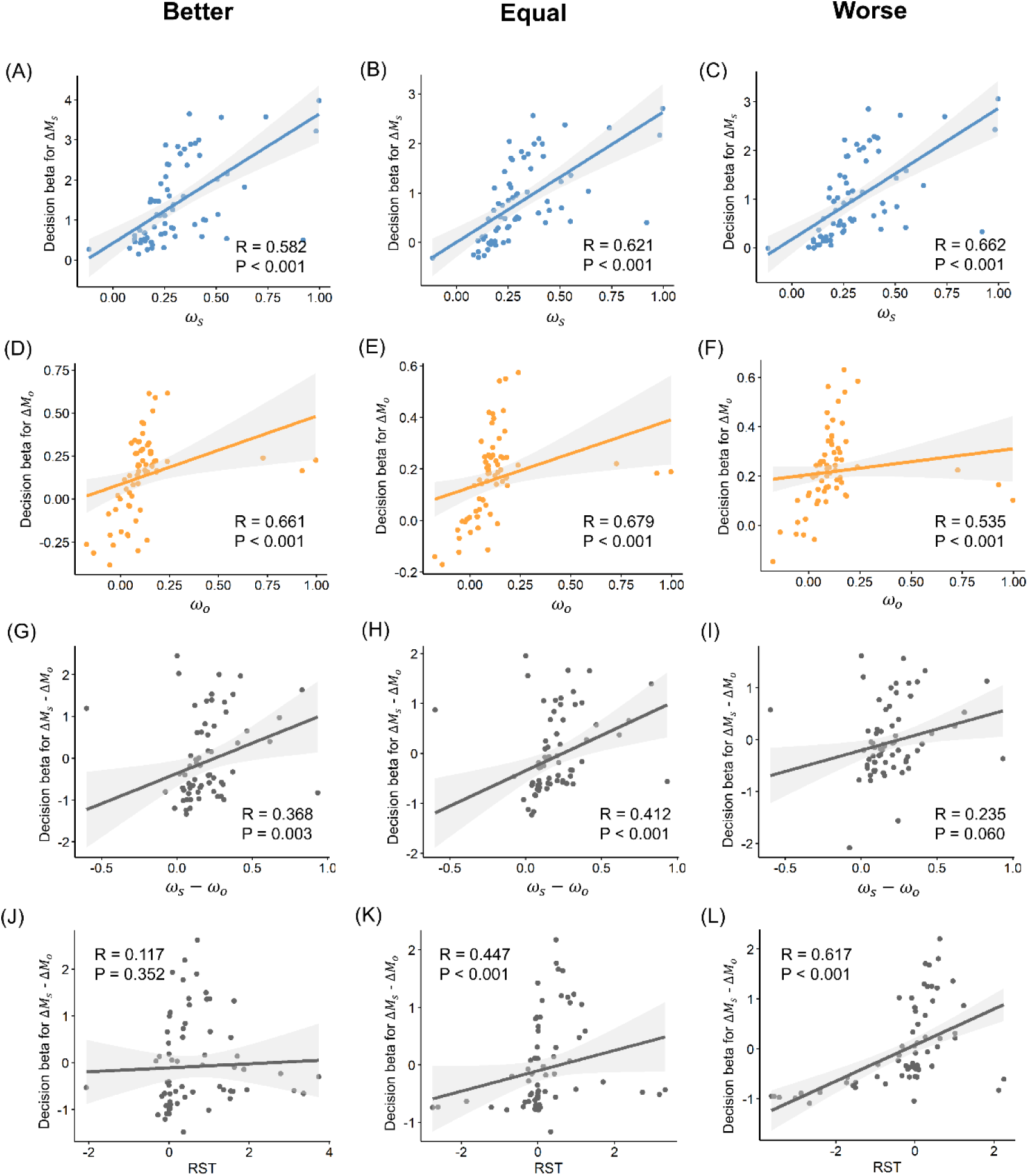
The correlation between model parameters and decision betas in study 1. (A-C), Robust correlation of drift weight and decision beta for Δ*M_s_* in better (A), equal (B) and worse (C) conditions. (D-F), Robust correlation of drift weight and decision beta for Δ*M_o_* in better (D), equal (E) and worse (F) conditions. (G-I), Robust correlation of relative drift weight and relative decision beta for Δ*M_s_* − Δ*M_o_* in better (G), equal (H) and worse (I) conditions. (J-L), Robust correlation of RST and relative decision beta (Δ*M_s_* − Δ*M_o_*) in better (J), equal (K) and worse (L) conditions.

**S6 Fig.**
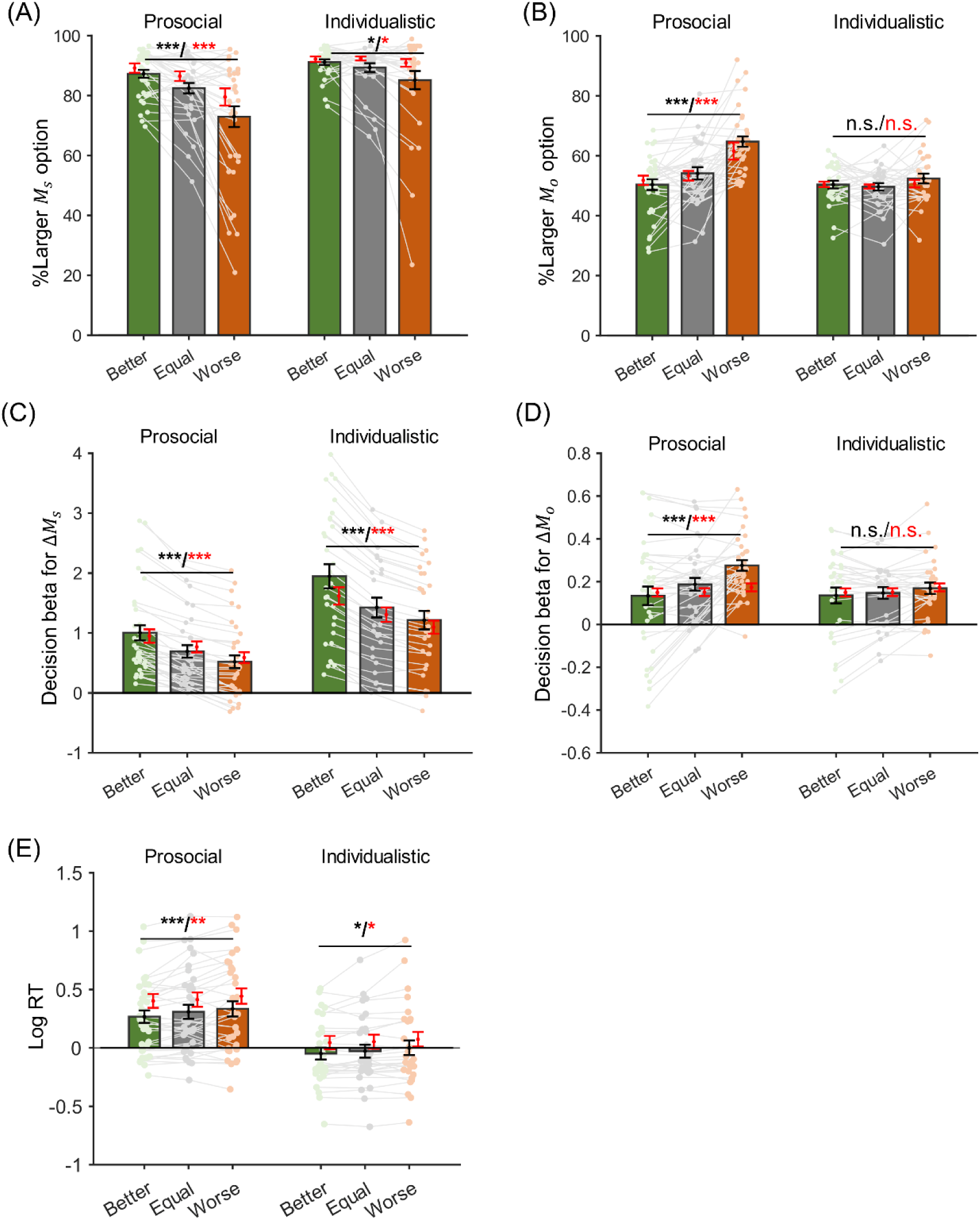
Behavioral differences between prosocial and individualistic subjects in study 1. (A) Proportion of larger *M_s_* option selection across social rank for the prosocial and individualistic groups. (B) Proportion of larger *M_o_* option selection across social rank for the prosocial and individualistic groups. (C) Decision betas of Δ*M_s_* across social rank for the prosocial and individualistic groups. (D) Decision betas of Δ*M_o_* across social rank for the prosocial and individualistic groups. (E) Mean log RT across social rank for the prosocial and individualistic groups. Asterisks across social rank denote significant main effect of one-way ANOVA. Asterisks across prosocial and individualistic groups denote significant interaction effect of two-way ANOVA (social rank×SVO group). Error bars represent s.e.m across subjects and red error bars represent model predictions from the cross-validation results (see methods). *** p < 0.001, ** p < 0.01 and * p < 0.05.

**S7 Fig.**
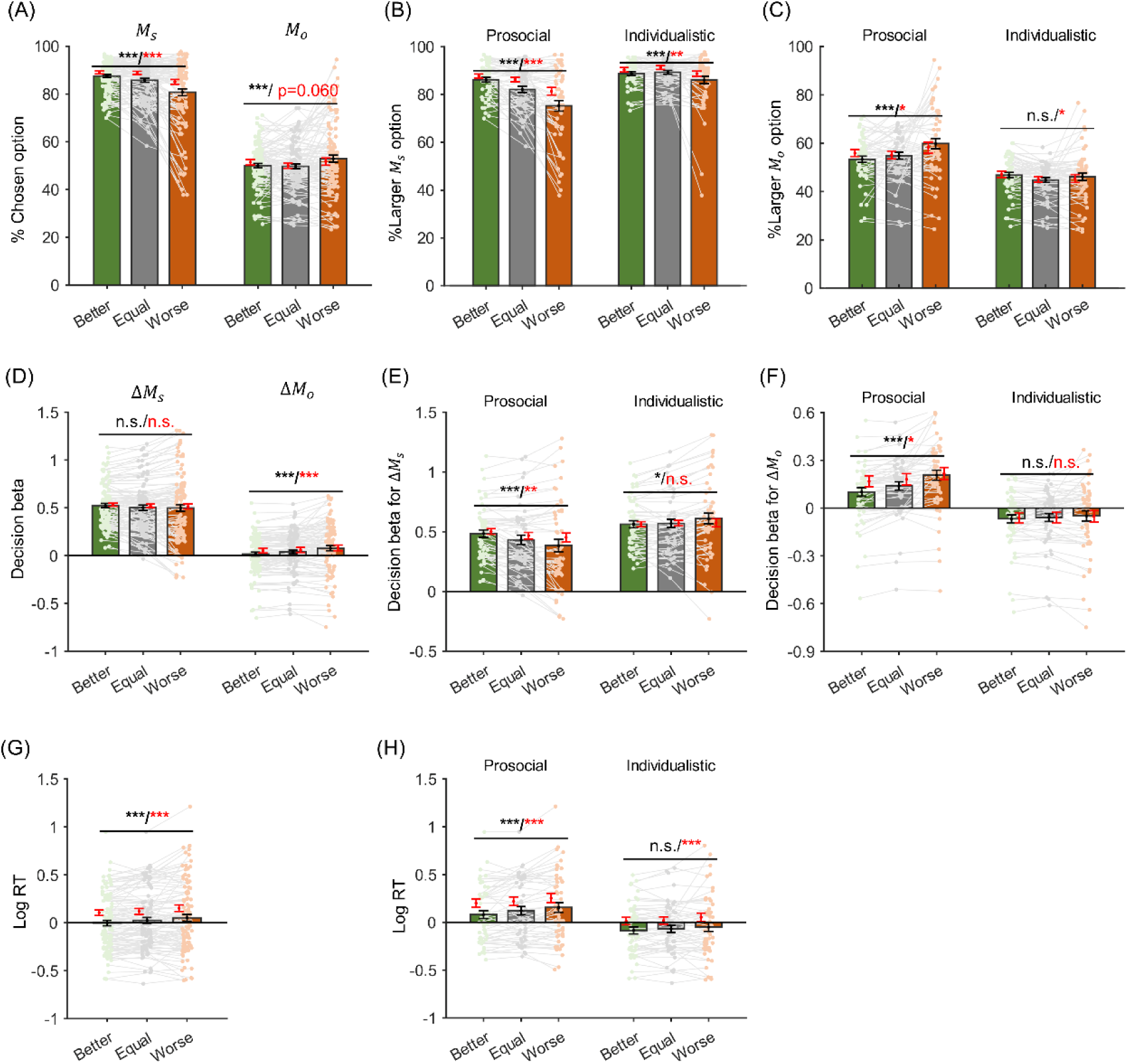
Behavioral results for the replication study. (A) Proportion of larger *M_s_* option selection across social rank for the prosocial and individualistic groups. (B) Proportion of larger *M_o_* option selection across social rank for the prosocial and individualistic groups. (C), Decision betas of Δ*M_s_* across social rank for the prosocial and individualistic groups., Decision betas of Δ*M_o_* across social rank for the prosocial and individualistic groups., Mean log RT across social rank for the prosocial and individualistic groups. Asterisks across social rank denote significant main effect of one-way ANOVA. Asterisks across prosocial and individualistic groups denote significant interaction effect of two-way ANOVA (social rank×SVO group). Error bars represent s.e.m across subjects and red error bars represent model predictions from the cross-validation results (see methods). *** p < 0.001, ** p < 0.01 and * p < 0.05.

**S8 Fig.**
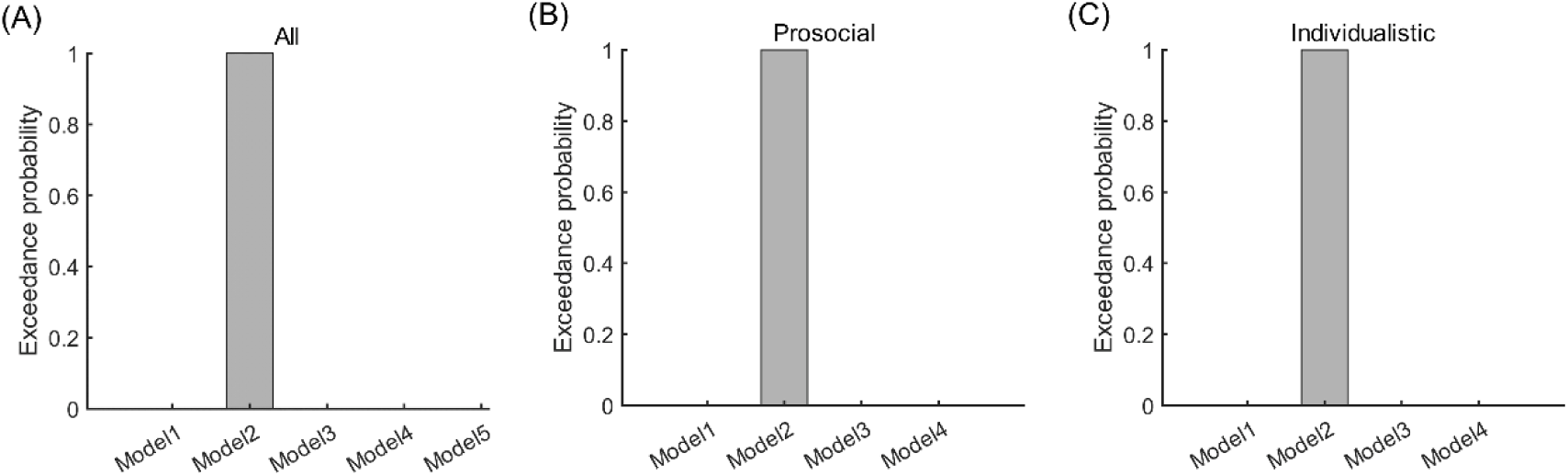
Bayesian model selection results for the replication study. Across all the subjects (A) or within the prosocial group (B) or individualistic group (C), model 2 (RST model) outperformed other models with the highest exceedance probability.

**S9 Fig.**
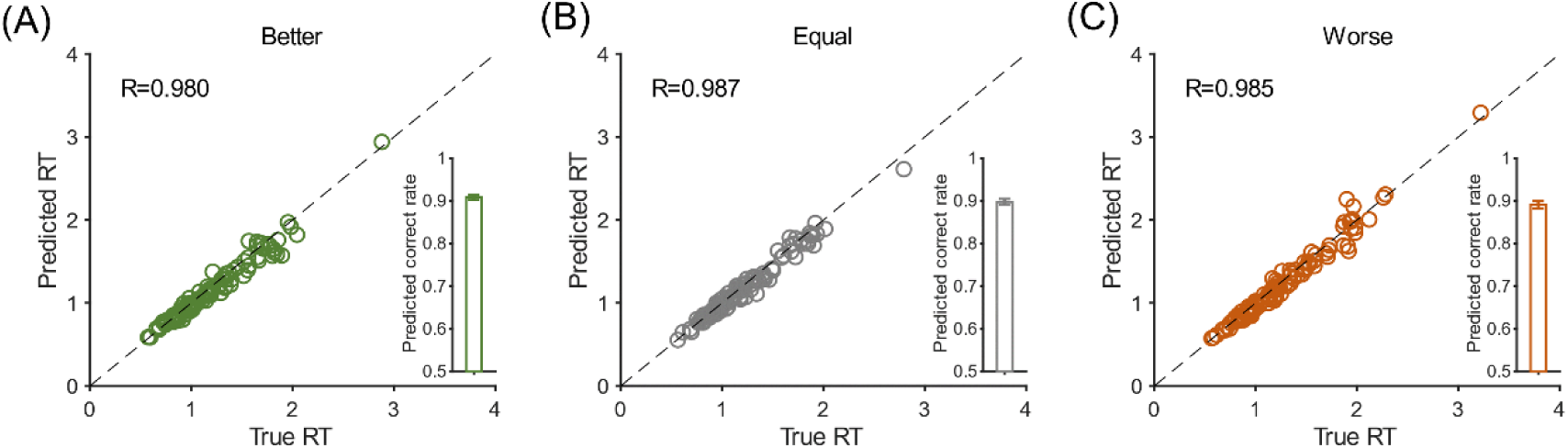
Cross-validation results for replication study. The correlation between predicted and participants’ actual RTs (empty circle) and choices (bar) across subjects in the better, equal and worse conditions.

**S10 Fig.**
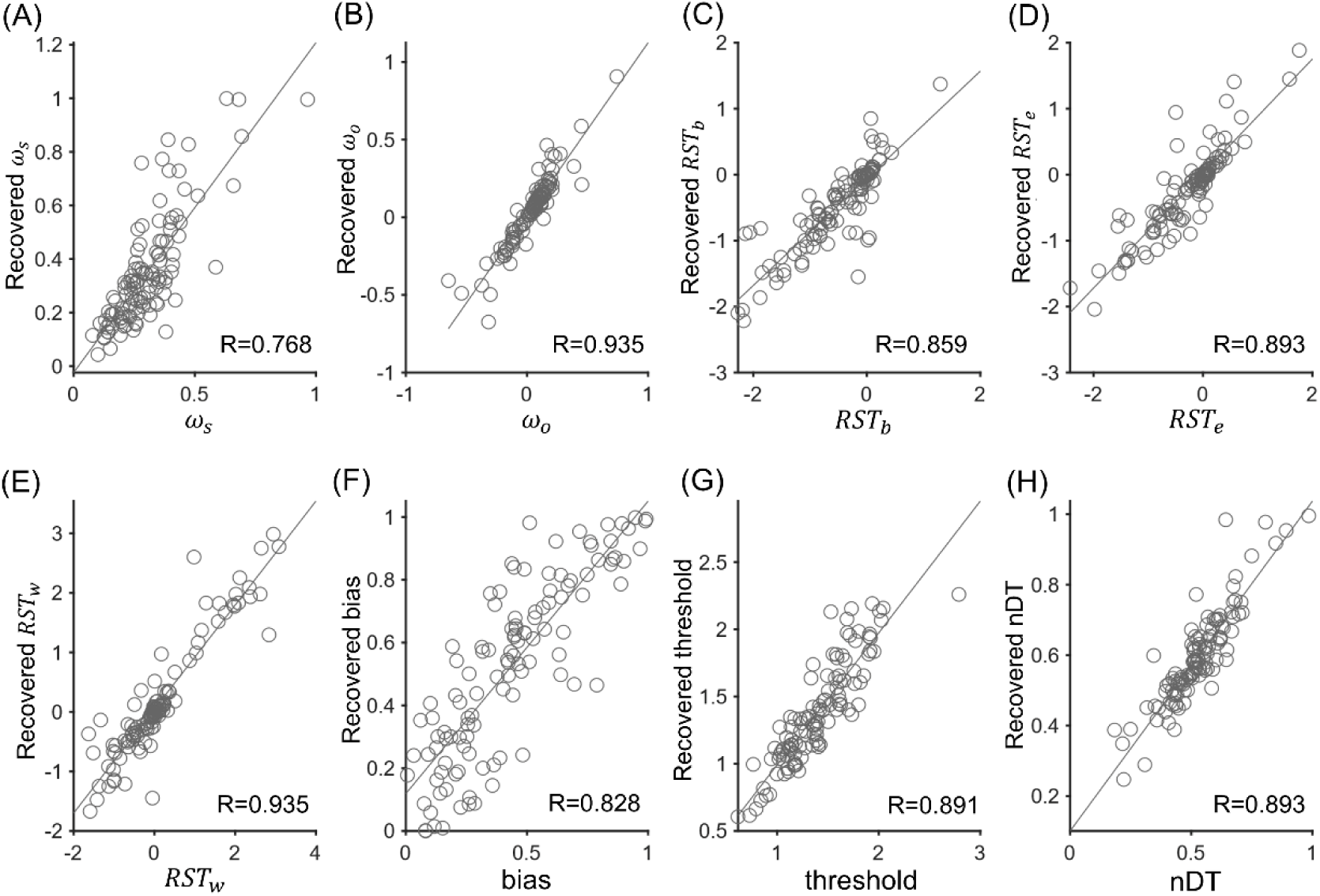
Parameter recovery analysis of the RST model for the replication study. Correlations between actual parameter and the model recovered parameters.

**S11 Fig.**
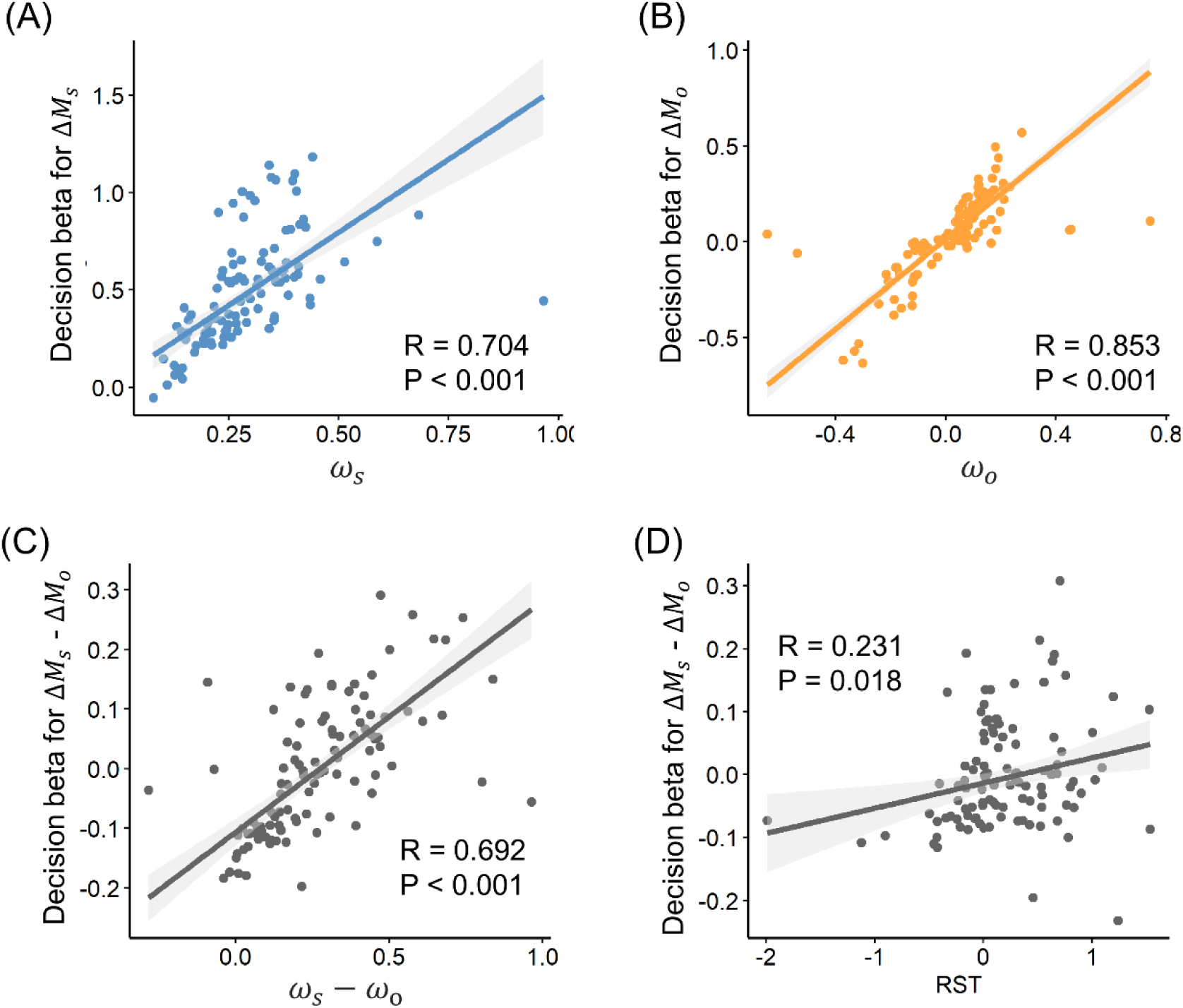
Correlations between model parameters and decision betas for the replication study. Robust correlation between drift weights and decision betas for Δ𝑀𝑠 (A) and Δ𝑀𝑜 (B) after controlling for the effect of RST. (C), Correlation between relative drift weights (Δ𝑀𝑠 - Δ𝑀𝑜) and relative decision betas (Δ𝑀𝑠 - Δ𝑀𝑜), controlling for the effect of RST. (D), Correlation between RSTs and relative decision betas, controlling for the effect of relative drift weight.

**S12 Fig.**
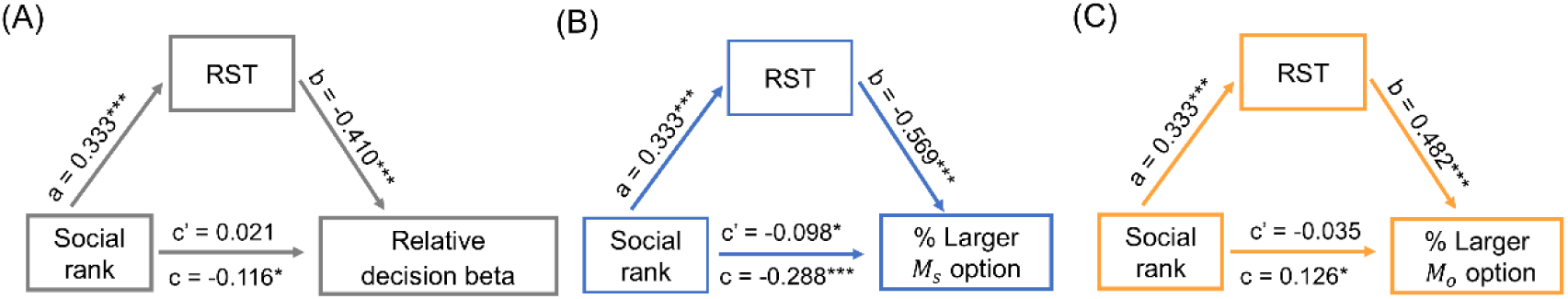
Mediation analysis for the replication study. (A), RST mediated the effect of social rank on subjects’ relative decision beta (Δ𝑀𝑠 - Δ𝑀𝑜). (B), RST partially mediated the effect of social rank on chosen larger 𝑀𝑠 proportions. (C), RST mediated the effect of social rank on chosen larger 𝑀𝑜 proportions.

**S13 Fig.**
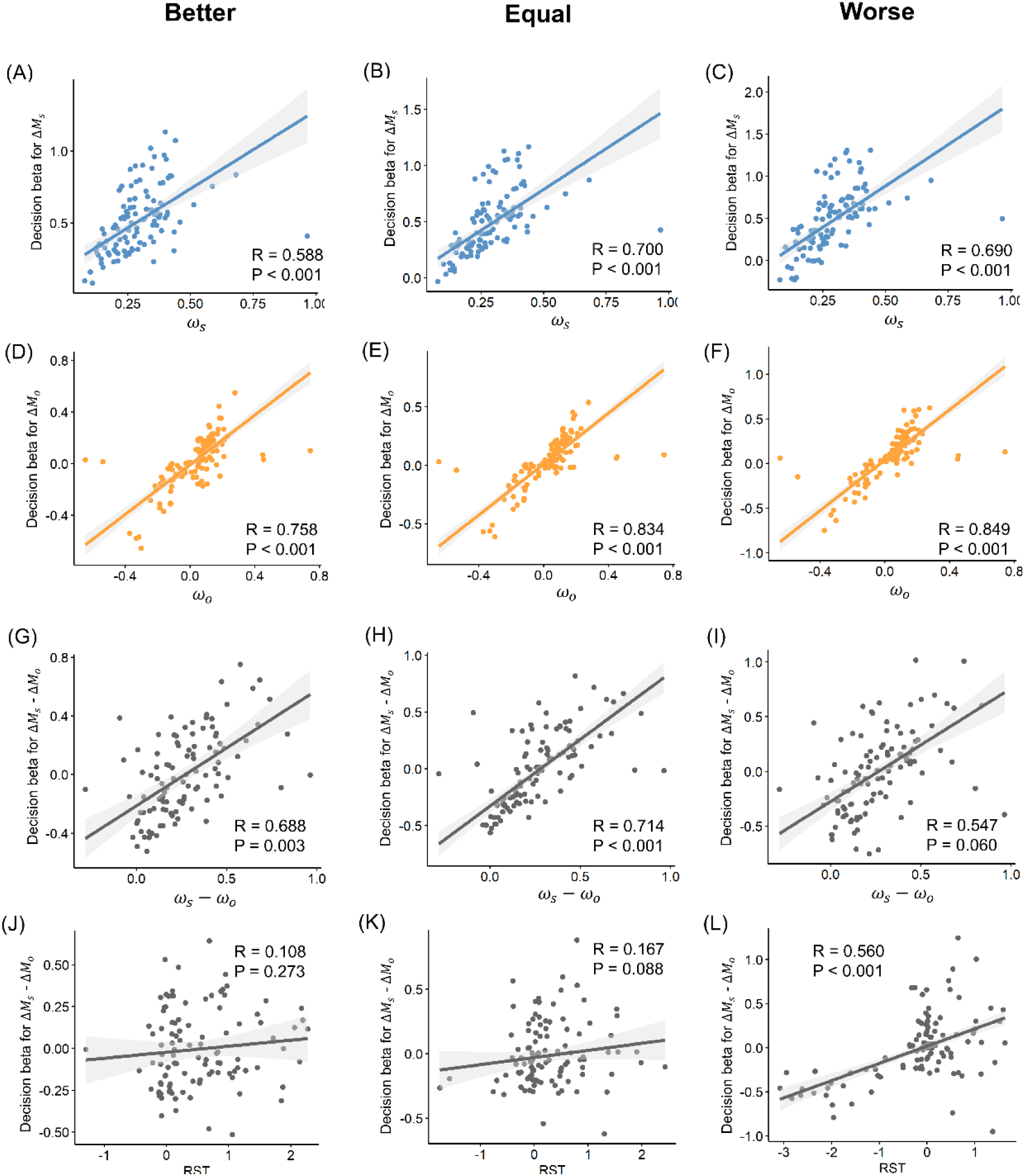
Model parameters correlation with decision regression coefficients for the replication study. (A-C), Robust correlation of drift weight and decision beta for Δ𝑀𝑠 in better (A), equal (B) and worse (C) conditions. (D-F), Robust correlation of drift weight and decision beta for Δ𝑀𝑜 in better (D), equal (E) and worse (F) conditions. (G-I), Robust correlation of relative drift weight and relative decision beta for Δ𝑀𝑠 − Δ𝑀𝑜 in better (G), equal (H) and worse (I) conditions. (J-L), Robust correlation of RST and relative decision beta (Δ𝑀𝑠 − Δ𝑀𝑜) in better (J), equal (K) and worse (L) conditions.

**S1 Table.** Specification and performance of candidate models. In the mtDDMs, 𝜔_𝑠_ and 𝜔_𝑜_ are the drift weights for Δ*M_s_* and Δ*M_o_*, RST is the relative start time of Δ*M_s_* over Δ*M_o_* . Bias: the initial starting point bias for choosing larger *M_s_* option in the evidence accumulation process. nDT: the non-decision time, which accounts for the extra amount of time required for subsequent motor action not related to evidence accumulation. Threshold: the evidence threshold or decision boundary for the decision.

| | $\omega_s$ | $\omega_o$ | RST | bias | nDT | threshold | Total<br>Parameters | BIC<br>(Study1) | BIC<br>(Replication study) |
| --- | --- | --- | --- | --- | --- | --- | --- | --- | --- |
| Model1 | 3 | 3 | 0 | 1 | 1 | 1 | 9 | 207223 | 337153 |
| Model2<br>(RST Model) | 1 | 1 | 3 | 1 | 1 | 1 | 8 | 207030 | 333708 |
| Model3 | 3 | 3 | 1 | 1 | 1 | 1 | 10 | 207375 | 337513 |
| Model4 | 3 | 3 | 3 | 1 | 1 | 1 | 12 | 208146 | 338731 |

**S2 Table.** Estimated model parameters for the winning model in study1 (Model 2, the RST model).

| Parameter | All<br>mean±s.d. | Prosocial<br>mean±s.d. | Individualistic<br>mean±s.d. |
| --- | --- | --- | --- |
| $\omega_s$ | 0.314±0.025 | 0.230±0.020 | 0.411±0.044 |
| $\omega_o$ | 0.120±0.023 | 0.103±0.011 | 0.140±0.049 |
| $RST_b$ | 0.618±0.120 | 0.793±0.202 | 0.414±0.101 |
| $RST_e$ | 0.179±0.121 | 0.227±0.199 | 0.123±0.125 |
| $RST_w$ | -0.343±0.155 | -0.613±0.232 | -0.029±0.186 |
| **bias** | 0.530±0.036 | 0.484±0.050 | 0.584±0.049 |
| **threshold** | 1.590±0.053 | 1.749±0.071 | 1.404±0.064 |
| **nDT** | 0.573±0.017 | 0.573±0.026 | 0.573±0.022 |

**S3 Table.** Estimated model parameters for the winning model in the replication study (Model 2, the RST model).

| Parameter | All<br>mean±s.d. | Prosocial<br>mean±s.d. | Individualistic<br>mean±s.d. |
| --- | --- | --- | --- |
| $\omega_s$ | 0.284±0.012 | 0.260±0.015 | 0.326±0.018 |
| $\omega_o$ | 0.037±0.018 | 0.076±0.020 | -0.017±0.028 |
| $RST_b$ | 0.496±0.063 | 0.510±0.109 | 0.521±0.076 |
| $RST_e$ | 0.294±0.059 | 0.080±0.095 | 0.459±0.079 |
| $RST_w$ | -0.195±0.093 | -0.553±0.148 | 0.187±0.118 |
| **bias** | 0.422±0.024 | 0.389±0.033 | 0.448±0.034 |
| **threshold** | 1.397±0.034 | 1.469±0.049 | 1.327±0.044 |
| **nDT** | 0.526±0.013 | 0.529±0.020 | 0.523±0.016 |

